# Computer Modeling of Bevacizumab Drug Distribution After Intravitreal Injection in Rabbit and Human Eyes

**DOI:** 10.1101/2023.05.05.539491

**Authors:** Jabia M. Chowdhury, Eduardo A. Chacin Ruiz, Matthew P. Ohr, Katelyn E. Swindle-Reilly, Ashlee N. Ford Versypt

**Author notes:** Corresponding author at 507 Furnas Hall, University at Buffalo, The State University of New York, Buffalo, NY 14260. Authors contributed equally (Jabia M. Chowdhury, Eduardo A. Chacin Ruiz).

## Abstract

Age-related macular degeneration (AMD) is a progressive eye disease that causes loss of central vision and has no cure. Wet AMD is the late neovascular form treated with vascular endothelial growth factor (VEGF) inhibitors. VEGF is the critical driver of wet AMD. One common off-label anti-VEGF drug used in AMD treatment is bevacizumab. Experimental efforts have been made to investigate the pharmacokinetic (PK) behavior of bevacizumab in vitreous and aqueous humor. Still, the quantitative effect of elimination routes and drug concentration in the macula are not well understood. In this work, we developed two spatial models representing rabbit and human vitreous to better understand the PK behavior of bevacizumab. This study explores different cases of drug elimination and the effects of injection location on drug concentration profiles. The models are validated by comparing them with experimental data. Our results suggest that anterior elimination is dominant for bevacizumab clearance from rabbit vitreous, whereas both anterior and posterior elimination have similar importance in drug clearance from the human vitreous. Furthermore, results indicate that drug injections closer to the posterior segment of the vitreous help maintain relevant drug concentrations for longer, improving bevacizumab duration of action in the vitreous. The rabbit and human models predict bevacizumab concentration in the vitreous and fovea, enhancing knowledge and understanding of wet AMD treatment.

## 1. Introduction

Retinal diseases are the most common causes of vision loss in all age groups. The retina is a multilayered nerve tissue that lines the back of the eye, and in its center lies the macula. The macula contains the photoreceptors responsible for central high-resolution color vision. A hypoxic condition in the macula can cause vascular endothelial growth factor (VEGF) overexpression. VEGF is a protein essential to blood vessel growth, and its overexpression at the macula promotes the formation of aberrant new blood vessels, causing the irremediable wet form of age-related macular degeneration (AMD). AMD damages the macula and affects the center of the visual field, limiting the patient’s ability to see clearly. AMD can cause irreversible central vision loss if it is not treated in time. Despite advances in treatment, AMD remains a significant public health issue and is the leading cause of severe vision loss in individuals older than 55 years in high-income countries ^1^.

In the United States, around 11 million people suffer from AMD ^2,3^, and the total is expected to double to 22 million by 2050^4^. AMD is generally found in people older than 65 years ^5,6^. There are two types of AMD: non-neovascular (dry) AMD and neovascular (wet) AMD. The most common AMD is dry AMD, which refers to the early stages of the disease when drusen deposition and possible geographic atrophy of the macula occur ^7^. Dry AMD can turn into wet AMD at any stage. Wet AMD is caused by abnormal blood vessel growth under the retina, and it causes severe vision loss by disrupting the normal anatomy of the retina ^8^. Early detection and treatment of AMD could reduce impairment of a patient’s vision.

Treatment of inner retinal diseases is challenging due to the anatomical barriers. Different ocular therapeutic approaches have been used to treat AMD, including intravitreal injection, drug implant, laser photocoagulation, and photodynamic therapy ^9^. Intravitreal injection of anti-VEGF drugs is considered the most common option in AMD treatment ^10^ since it allows high drug concentration to reach the retinal layer with minimal side effects from the localized ocular treatment, and the injection can be performed in-office. Bevacizumab (Avastin), a recombinant humanized monoclonal antibody that is one of the most readily available and commonly used anti-VEGF therapeutics, is considered in this work.

Some experimental studies have measured the pharmacokinetic (PK) profiles of bevacizumab in animal eye models ^11–15^ and in human eyes ^16,17^. Existing computational models of rabbit and human eyes have focused mainly on low molecular weight drugs ^18–27^ or on a multi-compartment approach to the eye ^28–32^. However, relatively less emphasis has been placed on developing a physiologically based three-dimensional (3D) PK model for the spatial distribution of bevacizumab in the eye.

In treating wet AMD, the therapeutic target is to control abnormal blood vessel growth in the macula. A region of particular interest in the macula is the fovea, located at the center and contains the largest concentration of cone photoreceptors that can be disrupted in AMD. Drug transport to this region depends on the vitreous properties. The vitreous has viscoelastic properties ^33–35^, comprises 80% of the eye’s volume ^35^, and is located between the lens and the retina. The rheological and mechanical properties of the vitreous change anisotropically with age ^35^. Here, we assume the vitreous to be homogeneous and constant, but our approach can incorporate spatial and temporal heterogeneity in properties in the future.

In this work, we develop 3D PK models for rabbit and human eyes with space and time dependence for intravitreal injection of bevacizumab in the vitreous. The vitreous is modeled by considering relevant anatomic parameters and routes of drug elimination. The models are validated using experimental PK results from rabbit and human eyes. The models allow for the calculation of bevacizumab concentration both in the vitreous and at the fovea. We determine the number of days VEGF is expected to be effective based on thresholds determined from *in vivo* and *in vitro* reference data.

## 2. Methods

### 2.1. Geometry

The 3D geometry used in this study is based on the anatomy and properties of the rabbit and human vitreous. The vitreous volume in rabbit eyes varies from 1.15 to 1.70 mL ^36^. The human vitreous is approximately three times bigger, and its volume varies between 4.22 and 5.43 mL ^37^. Our models use 1.44 and 4.79 mL volumes for the rabbit and human vitreous, respectively. Furthermore, the vitreous is considered semispherical in our model, influenced by the geometry from Khoobyar et al. ^38^; however, other physiologically based models of ocular geometry have been discussed elsewhere ^9^.

In the present models (Figure 1a), the vitreous is located in a 3D spherical domain with origin at (*x*_*v*_, *y*_*v*_, *z*_*v*_) = (0, 0, 0) mm and a radius *r*_*v*_ of 7.6 mm for rabbit vitreous and 10.9 mm for human vitreous. The sphere is truncated by cutting the anterior portion of the eye perpendicular to the *z*-axis at *z* = *z*_*h*_ = 4 mm and 7 mm for the rabbit and human eyes, respectively. This location represents the hyaloid membrane between the aqueous humor and the vitreous. The lens is considered as a smaller sphere of radius *r*_*l*_ at origin (*x*_*l*_, *y*_*l*_, *z*_*l*_) whose volume is subtracted from the vitreous domain. In the rabbit model, the center of the lens is located at (*x*_*l*_, *y*_*l*_, *z*_*l*_) = (0, 0, 4) mm, which corresponds to the truncation coordinate of the anterior portion of the rabbit eye in the *z* axis. However, in the human model, the lens origin is located at (*x*_*l*_, *y*_*l*_, *z*_*l*_) = (0, 0, 8) mm giving a 1 mm offset between the truncation plane in human and the lens origin. Together, the hyaloid membrane and the lens form the anterior boundary surfaces of the model domains. The vitreous semispherical outer surface is divided into two regions: anterior side and posterior side. The semispherical boundary defines the anterior vitreous outer surface for all *z* ≥ 0 and *r*_*v*_. The semispherical boundary defines the posterior vitreous outer surface for all *z <* 0 and *r*_*v*_. The fovea is at the outer surface of the vitreous at position (*x* _*f*_, *y* _*f*_, *z* _*f*_) = (0, 0, −*r*_*v*_), which is our target location for bevacizumab. Table 1 summarizes the geometric properties in our model.

**Table 1:** Model parameters used for the eye geometry, drug transport, Darcy’s law, and macular dimensions.

| Parameter <sup>Source</sup> | Rabbit | Human | Unit |
| --- | --- | --- | --- |
| Eye geometry |  |  |  |
| Vitreous origin $(x_v, y_v, z_v)$ | $(0, 0, 0)$ | $(0, 0, 0)$ | m |
| Vitreous radius $r_v$ | $7.6 \times 10^{-3}$ | $10.9 \times 10^{-3}$ | m |
| Vitreous volume | $1.44 \times 10^{-6}$ | $4.79 \times 10^{-6}$ | m<sup>3</sup> |
| Hyaloid membrane surface $(z_h)$ | $4 \times 10^{-3}$ | $7 \times 10^{-3}$ | m |
| Lens origin $(x_l, y_l, z_l)$ | $(0, 0, 4) \times 10^{-3}$ | $(0, 0, 8) \times 10^{-3}$ | m |
| Lens radius $r_l$ | $4.0 \times 10^{-3}$ | $4.9 \times 10^{-3}$ | m |
| Fovea location $(x_f, y_f, z_f = -r_v)$ | $(0, 0, -7.6) \times 10^{-3}$ | $(0, 0, -10.9) \times 10^{-3}$ | m |
| Injection bolus |  |  |  |
| Origin for anterior vitreous dose<br>$(x_a, y_a, z_a)$ | $(0, 4.9, -0.3) \times 10^{-3}$ | $(0, 6, 3) \times 10^{-3}$ | m |
| Origin for middle vitreous dose<br>$(x_m, y_m, z_m)$ | $(0, 2.5, -2.5) \times 10^{-3}$ | $(0, 3, -0.5) \times 10^{-3}$ | m |
| Origin for posterior vitreous dose<br>$(x_p, y_p, z_p)$ | $(0, 0, -4.5) \times 10^{-3}$ | $(0, 0, -4) \times 10^{-3}$ | m |
| Radius of intravitreal injection $r_i$ | $2.3 \times 10^{-3}$ | $2.3 \times 10^{-3}$ | m |
| Drug transport |  |  |  |
| Drug injection concentration $C_i$<br>Ref 11,13,39 | 0.167 | 0.167 | mol/m<sup>3</sup> |
| Drug diffusion coefficient $D$<br>Ref 28,40 | $1.2 \times 10^{-10}$ | $9.13 \times 10^{-11}$ | m<sup>2</sup>/s |
| Reaction rate $R$ <sup>Ref 41</sup> | 0 | 0 | mol/(m<sup>3</sup>·s) |
| Darcy’s law |  |  |  |
| Density $\rho$ <sup>Ref 42,43</sup> | 1000 | 1008 | kg/m<sup>3</sup> |
| Viscosity $\mu$ <sup>Ref 22</sup> | 0.001 <sup>a</sup> | 0.001 | Pa·s |
| Permeability $\kappa$ <sup>Ref 43</sup> | $5.8 \times 10^{-14}$ | - | m<sup>2</sup> |
| Permeability/viscosity $\kappa/\mu$<br>Ref 23,26,44 | - | $8.4 \times 10^{-11}$ | m<sup>2</sup>/(Pa·s) |
| Inlet $u$ at $z = z_l$ (fast convection)<br>Ref 38 | $1.5 \times 10^{-7}$ <sup>a</sup> | $1.5 \times 10^{-7}$ | m/s |
| Inlet $u$ at $z = z_l$ (slow convection) | $4.3 \times 10^{-8}$ | $3.4 \times 10^{-8}$ | m/s |
| Outflow $p$ at $r_v$ and $z < 0$ <sup>Ref 38</sup> | 0 <sup>a</sup> | 0 | Pa |
| Rabbit “macula” and human macula dimensions |  |  |  |
| Diameter <sup>Ref 45</sup> | $3.84 \times 10^{-3}$ <sup>b</sup> | $5.5 \times 10^{-3}$ | m |
| Volume <sup>Ref 46</sup> | $2.55 \times 10^{-9}$ <sup>b</sup> | $8.49 \times 10^{-9}$ | m<sup>3</sup> |

a: assumed same as human vitreous. b: values proportional to a human macula.

**Figure 1:**
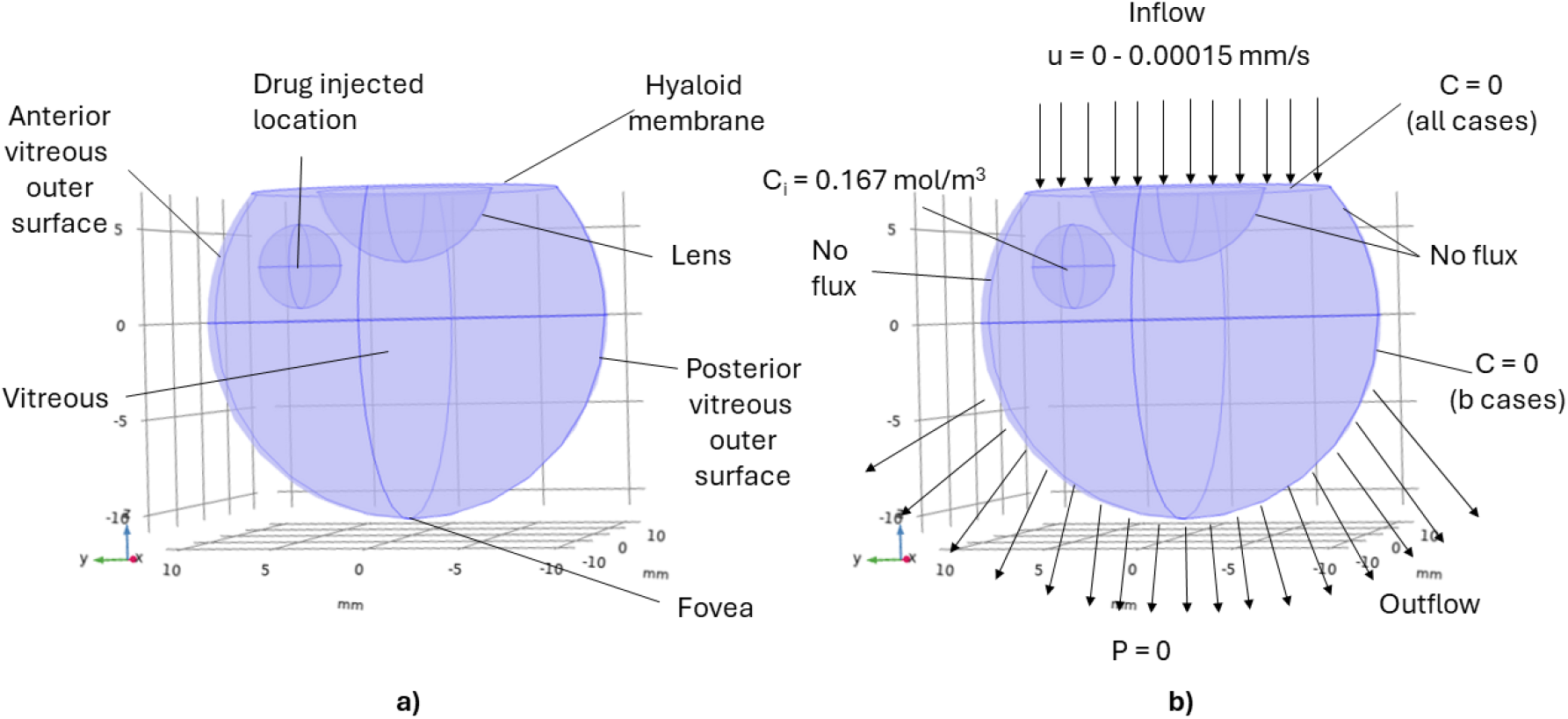
3D geometry of the human eye: a) anatomical region description and b) boundary conditions. The rabbit geometry considers the same anatomical regions and boundary conditions scaled to rabbit dimensions. Dimensions for both the rabbit and human geometries are in Table 1.

The injection is considered a sphere with a volume of 0.05 mL (Figure 1). Given that a 30G needle’s length usually varies between 12.7 to 15.75 mm ^47^, we consider different drug bolus origins to account for the potential variance in injection locations arising from differences in needle lengths and penetration depths. We define three injection locations: anterior vitreous, middle vitreous, and posterior vitreous (labeled as I, II, and III in Figure 2, respectively).

**Figure 2:**
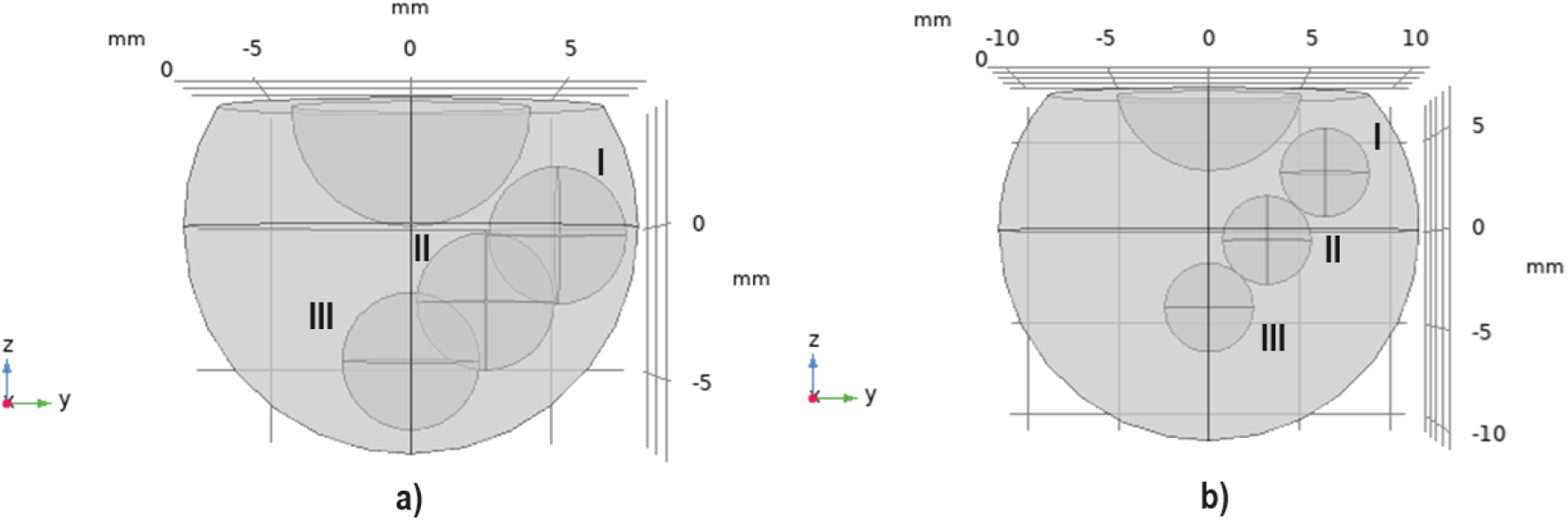
3D models for vitreous of a) rabbit and b) human. Drug intravitreal injection locations are labeled as I, II, and III for anterior, middle, and posterior locations, respectively.

### 2.2. Governing equations and parameters

#### 2.2.1. Drug transport

Our PK models capture the continuum of drug movement through the vitreous. The convection-diffusion-reaction equation governs drug transport in the vitreous:

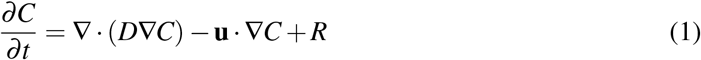

where *D* represents the diffusion coefficient of the drug, *C* is the drug concentration, **u** is the flow velocity, and *R* represents the reaction rate.

The PK models do not include the effect of metabolism and degradation of bevacizumab as both are negligible in the eye (*R* = 0) ^41^. The drug distribution rate depends on the drug diffusion coefficient value. The diffusion coefficients of bevacizumab in rabbit and human vitreous and drug initial concentration are tabulated in Table 1.

A typical dose concentration for bevacizumab administration is 1.25mg/0.05mL, which is used for both humans and rabbits ^11,13,39^. The concentration at the injected location is defined as *C*_*i*_ = 0.167 mol/m^3^. The initial drug concentration *C*_0_ in the domain is defined as:

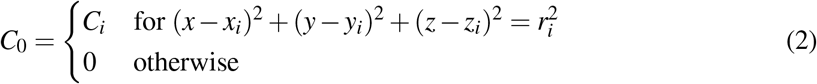

where *x*_*i*_, *y*_*i*_, and *z*_*i*_ are given in Table 1 for the following injection positions: *i* = *a* for the anterior of the vitreous nearest to the hyaloid membrane, *i* = *m* for the middle of the vitreous, and *i* = *p* for the posterior of the vitreous closest to the fovea. *r*_*i*_ is the resulting radius for a sphere of volume 0.05 mL.

#### 2.2.2. Fluid flow

Assuming the vitreous is a porous media, fluid flow in the vitreous can be modeled using Darcy’s law

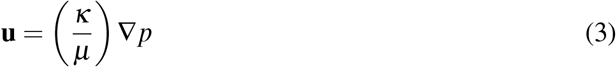

and the continuity equation

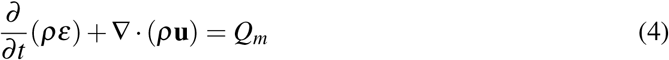

where **u** is the flow velocity, *κ* is the permeability of the vitreous, *µ* is the dynamic viscosity of the aqueous humor, *p* is the pressure, *ρ* is the density of the fluid, *ε* is the porosity, and *Q*_*m*_ is the mass source term, which is set to 0. The parameters used in Darcy’s law are tabulated in Table 1.

In our model, the vitreous is considered a porous structure with a porosity value of *ε* = 1 as in Penkova et al. ^48^. A reference pressure of *p* = 1 atm was used. Darcy’s law’s initial condition was zero throughout the domain (*p* = 0) before solving the steady-state pressure distribution pre-injection.

Slow (8×10^*−*9^ m/s) and fast (1.5×10^*−*7^ m/s) inlet velocities to the vitreous through the hyaloid membrane have been previously considered from Heljak and Swieszkowski ^49,50^ for the slow case and Khoobyar et al. ^38^ for the fast case. We calculate a slow inlet velocity based on a constant aqueous humor production rate of 3 µL/min. Assuming that 7% of the aqueous humor production enters the rabbit vitreous and 10% enters the human vitreous ^26^ and using the hyaloid membrane surface areas of 0.81 cm^2^ for rabbits and 1.47 cm^2^ for humans (from our geometric models), we determine the corresponding slow inlet velocities. We use the same value as Khoobyar et al. ^38^ for the fast inlet.

Furthermore, a recent study ^51^ demonstrated that fluid velocity generated from eye globe expansion and deflation after intravitreal drug injections has a negligible effect on drug distribution in the eye; therefore, we do not consider any vitreous volume changes in our model.

### 2.3. Boundary conditions

The major routes of drug elimination in the eye are through the aqueous humor drainage (anterior elimination route) and blood-retinal barrier (posterior elimination route), with the anterior elimination route being the dominant one ^41,43^. We consider the elimination routes via the boundary conditions (Figure 1b). We only consider the extreme cases of zero concentration (*C* = 0, which is equivalent to infinitely fast flux from the boundary) and no flux; however, others have considered the boundaries to be membranes with mass transport limitations (e.g., finite flux) between tissues ^28–32,38,52^.

The no-flux boundary condition for drug mass transport is defined by

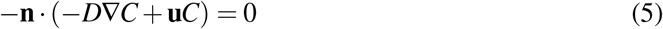

where **n** is the unit normal vector, *D* is the the diffusion coefficient of the drug, *C* is the drug concentration, and **u** is the fluid velocity.

The no-flux boundary condition for fluid flow is defined by

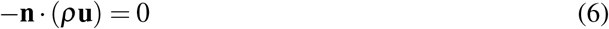

where **n** is the unit normal vector, *ρ* is the density, and **u** is the fluid velocity.

#### 2.3.1. Anterior elimination: hyaloid membrane

We consider the hyaloid membrane surface as the boundary for the anterior elimination route. We define a boundary condition of *C* = 0 at the hyaloid membrane’s surface based on the idea that the aqueous humor mass transfer rate is fast compared to the rate at which the drug reaches the anterior segment. Therefore, the drug is cleared as soon as it reaches the anterior segment. Moreover, a portion of the aqueous can permeate through the vitreous ^23^. Our models consider this to be an input velocity through the hyaloid membrane.

#### 2.3.2. Impermeable surfaces: Lens and anterior portion of the vitreous outer surface

The lens and the anterior portion of the vitreous outer surface (*z* ≥ 0 and *r*_*v*_) are assumed to be impervious to fluids and drugs. Therefore, no-flux boundary conditions (Equations (5) and (6)) are applied at these tissues implying that there is no velocity (*u* = 0) across these boundaries, and no drug can leave the lens and anterior portion of the vitreous outer surface.

#### 2.3.3. Posterior elimination: Posterior portion of the vitreous outer surface

We consider the posterior portion of the vitreous outer surface (*z <* 0 and *r*_*v*_) as the boundary for the posterior elimination route. Given that the blood can be regarded as a perfect sink and that the retinal layer surrounding this tissue is thin, a concentration of *C* = 0 is considered for cases (described in Section 2.4) with posterior elimination. Furthermore, a reference pressure of *p* = 0 Pa is set as the outflow boundary condition for the fluid flow (Figure 1b). No-flux boundary conditions are used for cases without posterior elimination.

### 2.4. Case studies

We investigate four cases studies with our models to explore different combinations of convection and drug elimination routes (Table 2). The vitreous is modeled as a porous viscoelastic material for the cases where convection is included (cases denoted by “1”). The aqueous humor fluid produced from the ciliary body permeates the vitreous through the hyaloid membrane, and we consider it as an inlet fluid velocity in our models following another published model that considered convection ^38^ (Figure 1b). Other models have also considered convection in the vitreous and point to the importance of this transport mechanism for slow diffusing macromolecules ^26^. Slow convection is considered for the three injection locations studied. Another alternative common assumption is that there is no fluid flow in the vitreous when modeling transport across the vitreous ^24,53^. In the cases denoted by “2”, we consider zero inlet velocity (*u* = 0) in the vitreous, meaning there is no convection and only diffusion across the vitreous. Fast convection is only considered at the middle vitreous position as a case study on the effect of convection on drug distribution in the eye. As most previous studies have considered slow convection or no convection, we only consider fast convection from one reference ^38^ in injection location rather than repeating the analysis for all the positions. The resulting flow from Equations (3) and (4) at steady-state is coupled to the convection-diffusion equation (Equation (1)) for the drug transport.

**Table 2:** Summary of case studies.

| | Convection | Anterior elimination ($C = 0$) | Posterior elimination ($C = 0$) |
| --- | --- | --- | --- |
| Case 1a | ✓ | ✓ | X |
| Case 1b | ✓ | ✓ | ✓ |
| Case 2a | X | ✓ | X |
| Case 2b | X | ✓ | ✓ |

The mechanism for drug clearance in the eye is not fully understood. High molecular weight compounds are usually eliminated via the anterior route ^10,31,54^. A recent study observed the elimination routes of anti-VEGF therapeutics in both human and rabbit vitreous ^41^. The results suggested that the anterior elimination route is dominant compared to the posterior route for bevacizumab and other anti-VEGF drugs in both human and rabbit eyes. This dominance also depends on the eye’s aqueous humor flow rate. However, monoclonal antibodies such as bevacizumab are lipophilic compounds ^55,56^. Lipophilicity allows molecules to cross the retina and be eliminated through the posterior route ^10^.

In our simulations, we consider two elimination routes. Only anterior elimination is considered in the cases denoted by “a” (*C* = 0 at the hyaloid membrane surface and no-flux boundary conditions on all the other boundaries). In the cases denoted by “b”, both anterior and posterior elimination (*C* = 0 boundary conditions at the hyaloid membrane and the posterior portion of the vitreous outer surface) are considered. We do not consider a posterior elimination route alone, given that experimental results point towards the prevalence of the anterior elimination route.

We combine the two cases for convective flow modeling in the vitreous with the two elimination cases for a total of four case studies (Table 2). We study convection in the vitreous with anterior elimination only (Case 1a), convection with both anterior and posterior elimination (Case 1b), no convection with anterior elimination only (Case 2a), and no convection flow with both anterior and posterior elimination (Case 2b).

### 2.5. Solution methods and meshing

The rabbit and human eye geometries were created, and model equations were formulated and solved in COMSOL Multiphysics 6.1^57^. The Transport of Diluted Species interface was used for the mass transport of bevacizumab. The Darcy’s Law interface was used for fluid flow through the vitreous.

The mesh was generated using the “extra fine” physics-controlled mesh option in COMSOL. The rabbit model consisted of 162,750 elements and 7,252 boundary elements, and the human model consisted of 185,119 elements and 7,076 boundary elements. A mesh independence test was performed using drug concentration results at different mesh sizes. The results were determined to be independent of the mesh settings (Figures S1 and S2).

### 2.6. Concentration and half-life calculations

From our 3D PK model, we calculate drug concentration in the vitreous and at the fovea for the different case studies described. Concentration in the vitreous is calculated as the volume average in the vitreous domain for both the rabbit and human models using a volume probe in COMSOL.

Bevacizumab concentration at the fovea for Case 1a and Case 2a is calculated using a point probe at the locations specified as (*x* _*f*_, *y* _*f*_, *z* _*f*_) (Table 1). This method cannot be applied for Cases 1b and 2b since *C* = 0 is an established boundary condition in these cases. Therefore, we determine the bevacizumab flux using a point probe in the same locations (*x* _*f*_, *y* _*f*_, *z* _*f*_). Then, we use the diameter and volumetric information of the macula (Table 1) to approximate its surface area. Surface area multiplied by flux gives an amount per time, which is then divided by the macula volume for a concentration per time. We then calculate a rolling daily average concentration assuming that the fovea is just on the outside of the point on the vitreous boundary (Figures S3–S6).

There is no clearly defined macula in the rabbit eye ^58^; rabbits have a horizontal area with higher cell density called a visual streak ^59^. However, we are interested in comparing the amount of drug leaving the vitreous and reaching the retina near the macula region in both species, so we assumed a rabbit “macula” proportional to the scale of a human macula. Table 1 contains the parameters used for the human macula and the rabbit macula approximation. The center point of the macula is taken to be the fovea in both species.

An injection of 12.5 µg of bevacizumab was shown to partially or completely reduce leakage of neovascularization at 7 days after dosing in patients with proliferative diabetic retinopathy ^60^. Dividing the 12.5 µg of bevacizumab over the human vitreous volume in our model provides a conservative *in vivo* threshold vitreous concentration of 2.6 µg/mL. Therefore, after calculating the bevacizumab concentration at the vitreous with each model, the results determine the number of days when the bevacizumab concentration is above this threshold, which is defined as “duration of action”. Furthermore, after 7 days of a 12.5 µg intravitreal bevacizumab injection, the concentration at the human fovea was calculated to be 0.44 µg/mL according to our model. This is consistent with the *in vitro* minimum bevacizumab concentration of 500 ng/mL (0.5 µg/mL) that was reported to inhibit VEGF completely ^61^. The value of 0.5 µg/mL was considered as an *in vitro* threshold concentration for the macula concentrations due to the good agreement with our calculated value.

Half-lives were calculated by interpolating concentration results. Starting from an arbitrary base concentration at time points of 3 days for rabbits and 5 days for humans, the time at which the concentration reduced by half was obtained through interpolation of the concentration results.

### 2.7. Validation

WebPlotDigitizer ^62^ was used to extract data from plots in previously published experimental studies on rabbit and human eyes to validate the model. These experiments measured bevacizumab concentration in the vitreous, but none measured the drug concentration at the macula in human eyes or at the visual streak in rabbit eyes. The experiments followed a similar methodology. Briefly, 1.25 mg/0.05 mL of bevacizumab was injected 1–4 mm behind the surgical limbus in the vitreous supertemporal quadrant using 30G needles. Model simulations were performed under the same conditions as these experiments.

### 2.8. Code availability

We have provided the COMSOL codes, results spreadsheets, and analysis scripts for generating plots in a repository at https://github.com/ashleefv/PBPKHumanRabbitEyes^63^.

## 3. Results

### 3.1. Effects of injection location on bevacizumab distribution after intravitreal injection

The calculated bevacizumab vitreous half-lives for the simulated cases at each injection location are tabulated in Table 3. Figure 3 shows the normalized vitreous concentration profiles of bevacizumab when the injection dose is given at the anterior (a), middle (b), and posterior (c) of the rabbit vitreous and at the anterior (d), middle (e), and posterior (f) of the human vitreous. Figure S7 shows the same concentration profiles of bevacizumab as Figure 3 except on a logarithmic scale, so that the later timepoints are more visible. Regardless of injection location, the slowest clearance in the vitreous happens when considering no convection and anterior elimination only (Case 2a). The fastest clearance from the eye is obtained when considering both anterior and posterior elimination with or without convection (Case 1b and Case 2b, respectively). Intermediate clearance and bevacizumab concentrations are obtained when considering convection through the vitreous and anterior elimination (Case 1a). Spatial distribution plots of bevacizumab concentration at different times in the human vitreous are available for each case in Figures S8–S11.

**Table 3:** Calculated bevacizumab vitreous half-lives in days.

| Injection location | Case 1a | Case 1b | Case 2a | Case 2b |
| --- | --- | --- | --- | --- |
| Rabbit |  |  |  |  |
| Anterior vitreous | 2.12 | 0.38 | 3.26 | 0.38 |
| Middle vitreous (slow convection) | 2.12 | 0.38 | 3.27 | 0.38 |
| Middle vitreous (fast convection) | 1.00 | 0.32 | 3.27 | 0.38 |
| Posterior vitreous | 2.14 | 0.38 | 3.27 | 0.38 |
| Human |  |  |  |  |
| Anterior vitreous | 5.29 | 1.20 | 10.00 | 1.27 |
| Middle vitreous (slow convection) | 5.36 | 1.19 | 10.25 | 1.27 |
| Middle vitreous (fast convection) | 1.62 | 0.68 | 10.25 | 1.27 |
| Posterior vitreous | 5.46 | 1.19 | 10.56 | 1.27 |

**Figure 3:**
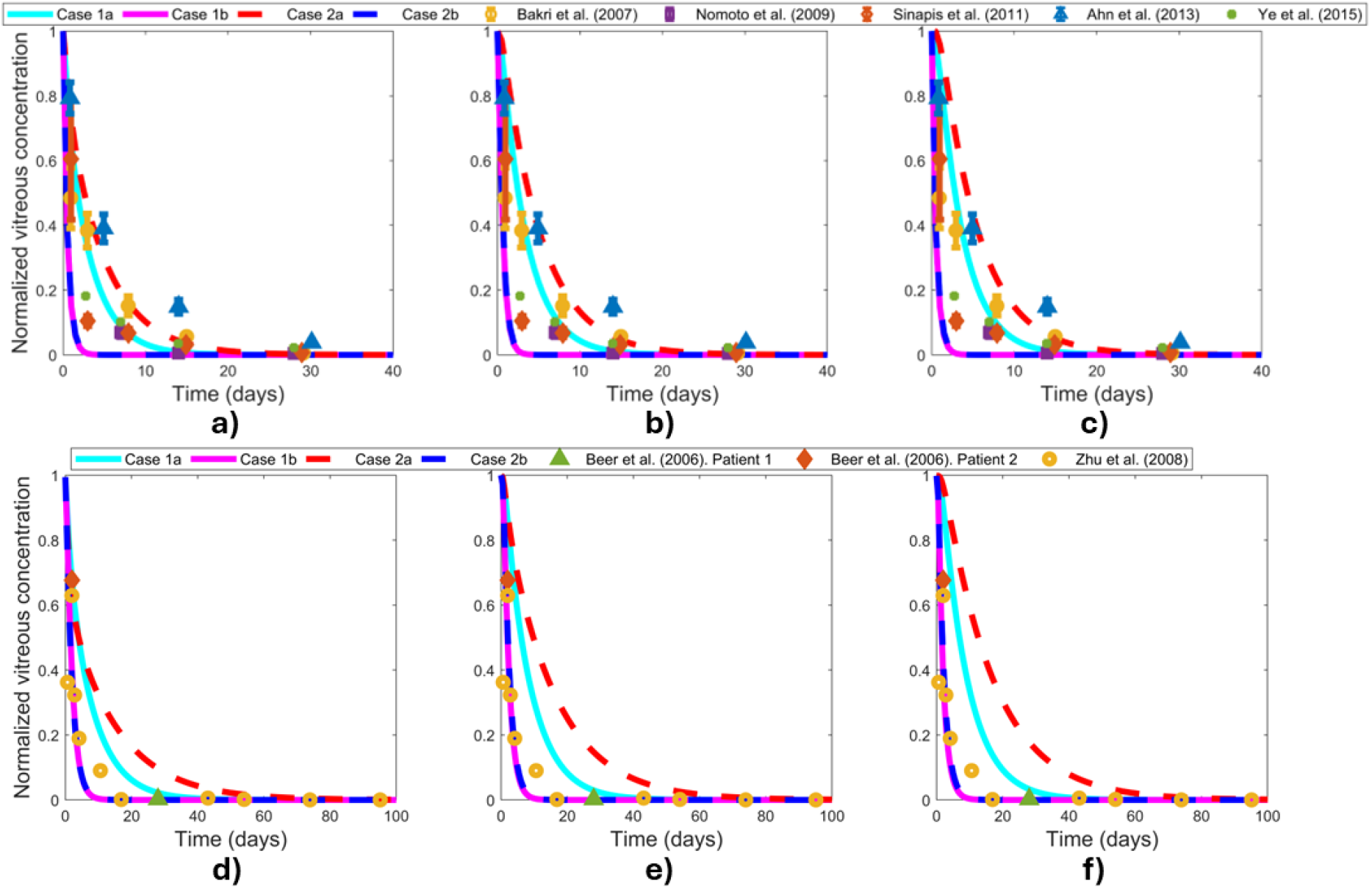
Normalized vitreous concentration in the rabbit and human eye. Injection locations: a) rabbit anterior vitreous, b) rabbit middle vitreous, c) rabbit posterior vitreous, d) human anterior vitreous, e) human middle vitreous, and f) human posterior vitreous. Case 1a: slow convective flow with anterior elimination only, Case 1b: slow convective flow with both anterior and posterior elimination, Case 2a: no convective flow with anterior elimination, and Case 2b: no convective flow with both anterior and posterior elimination. Model results for each case are shown as curves. Experimental data (markers) from Bakri et al. ^11^, Nomoto et al. ^12^, Sinapis et al. ^13^, Ahn et al. ^14^, Ye et al. ^15^, Beer et al. ^16^, and Zhu et al. ^17^.

A minimum concentration of 2.6 µg/mL is likely required to completely prevent angiogenesis (see Section 2.6 for the rationale behind this *in vivo* threshold vitreous concentration). Table 4 summarizes the predicted bevacizumab duration of action for the study cases at the different injection locations. Considering the effects of injection location only for both Case 1b and Case 2b, our rabbit model predicts that the concentration of bevacizumab goes below the threshold after 3 days, independent of the injection position. In Case 1a, bevacizumab concentration above the threshold is maintained in the rabbit vitreous for 17–18 days for injections located at the anterior, middle, and posterior vitreous, respectively. Case 2a maintains a duration of action of bevacizumab for 26–28 days in rabbits, depending on the injection location. In humans, Case 1b bevacizumab concentration goes below the 2.6 µg/mL threshold after 8–8.5 days. Similarly, the duration of action for Case 2b for humans varies in the range of 8.5–9 days. For Case 1a in humans, the predicted duration of action is in the range of 34–37 days. Case 2a predicts a duration of action longer than two months in humans.

**Table 4:** Calculated bevacizumab duration of action in the vitreous in days.

| Injection location | Case 1a | Case 1b | Case 2a | Case 2b |
| --- | --- | --- | --- | --- |
| Rabbit |  |  |  |  |
| Anterior vitreous | 17.0 | 3.0 | 26.0 | 3.0 |
| Middle vitreous (slow convection) | 18.0 | 3.0 | 27.5 | 3.0 |
| Middle vitreous (fast convection) | 8.5 | 2.5 | 27.5 | 3.0 |
| Posterior vitreous | 18.0 | 3.0 | 28.0 | 3.0 |
| Human |  |  |  |  |
| Anterior vitreous | 34.0 | 8.0 | 61.5 | 8.5 |
| Middle vitreous (slow convection) | 36.0 | 8.5 | 67.5 | 9.0 |
| Middle vitreous (fast convection) | 11.5 | 5.0 | 67.5 | 9.0 |
| Posterior vitreous | 37.0 | 8.0 | 70.5 | 8.5 |

Figure 4 shows bevacizumab concentration in the rabbit “fovea” when the injection dose is given at the anterior (a), middle (b), and posterior (c) of the rabbit vitreous and in the human fovea for injections given at the anterior (d), middle (e), and posterior (f) of the human vitreous. The results are consistent with the bevacizumab concentration profiles in the vitreous obtained in Figure 3. When posterior elimination is considered (Case 1b and Case 2b) for all drug locations, a rapid initial increase in the bevacizumab concentration at the fovea is observed, followed by a sharp decrease. When posterior elimination is not considered (Case 1a and Case 2a), the maximum concentration achieved at the fovea is lower, but clearance is also slower. The location of the dose being closest to the fovea (moving closer is shown from left to right across Figure 4) yields the highest concentrations of bevacizumab delivered to the fovea (Figure 4c, f).

**Figure 4:**
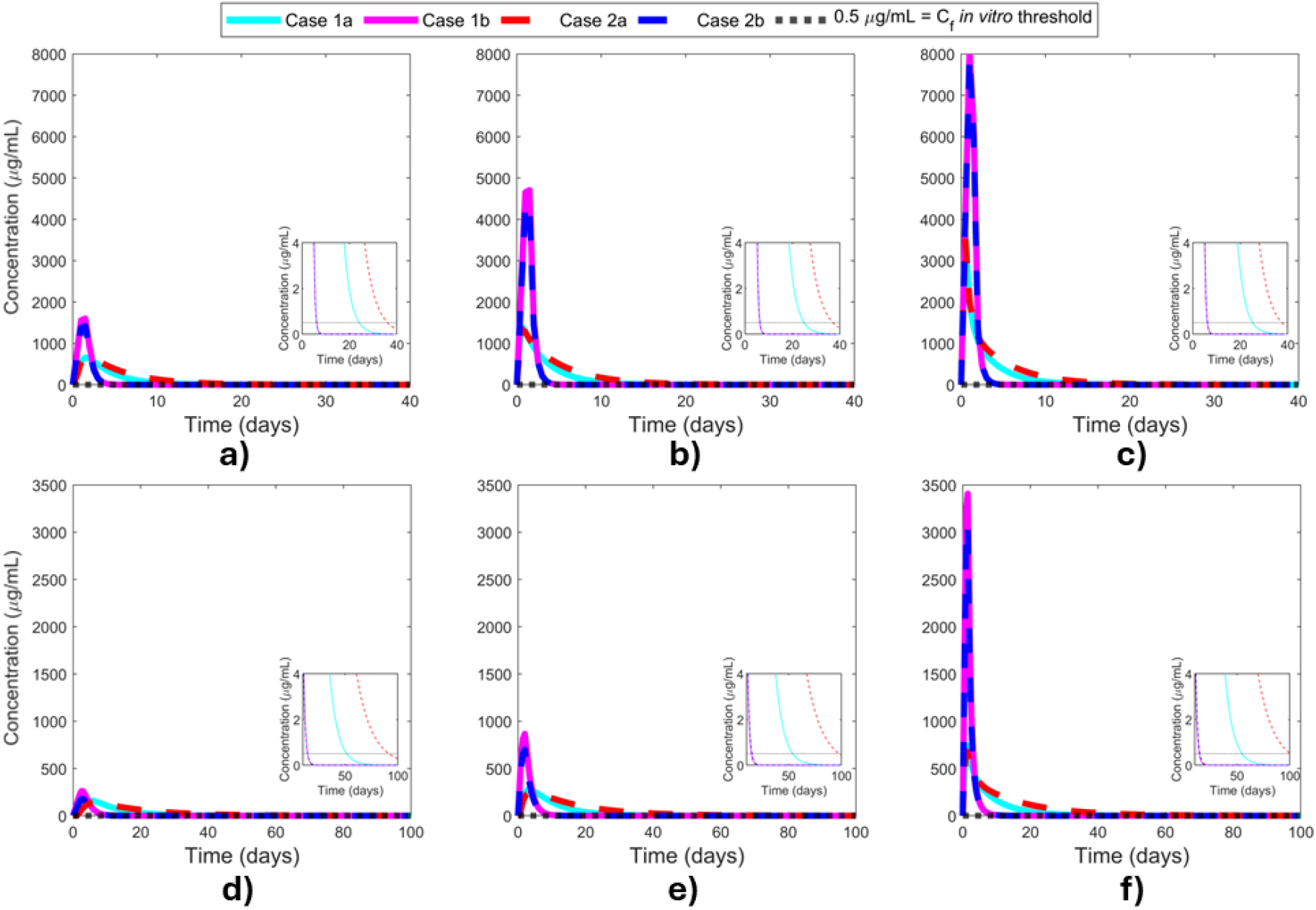
Drug concentration at the rabbit “fovea” and human fovea. Injection locations: a) rabbit anterior vitreous, b) rabbit middle vitreous, c) rabbit posterior vitreous, d) human anterior vitreous, e) human middle vitreous, and f) human posterior vitreous. Case 1a: slow convective flow with anterior elimination only, Case 1b: slow convective flow with both anterior and posterior elimination, Case 2a: no convective flow with anterior elimination, and Case 2b: no convective flow with both anterior and posterior elimination. *C*_*f*_ : *in vitro* threshold concentration at the fovea.

### 3.2. Effects of convection on bevacizumab distribution after intravitreal injection in the middle vitreous

We compared the cases of slow and fast aqueous humor inlet at the rabbit and human eyes when the injection was placed in the middle vitreous. The velocity profiles are shown in Figure S12. Using the model, we calculated the spatial distribution of pressure in the human vitreous domain and found a small pressure drop of around 2 Pa for slow convection cases (Figure S13a) and below 10 Pa for fast convection cases (Figure S13b).

Figure S14 shows the normalized concentration profiles of the drug bevacizumab in the rabbit vitreous when the injection dose is given at the middle vitreous for slow (a) and fast (b) convection and in the human vitreous when the injection dose is given at the middle vitreous for slow (c) and fast (d) convection. The cases of interest in Figures S14 and S15 are Case 1a and Case 1b, since Case 2a and Case 2b do not consider convection and, therefore, present no or subtle differences between the plots. The effects of the fast convection are most clearly observable for Case 1 in Figure S14 as the corresponding curves shift markedly between the plots. In Figure S15 the early intensity at the fovea is different between all four cases and in each convection scenario plotted. The time above the *in vitro* threshold concentration at the fovea changes most dramatically for Case 1a (Figure S15 insets). Bevacizumab vitreous half-lives for the cases of slow and fast convection are reported in Table 3. The duration of action of bevacizumab is tabulated in Table 4. The bevacizumab concentration profiles in the rabbit “fovea” and the human fovea are shown in Figure S15 for the different convection scenarios.

A faster aqueous humor inlet reduces the duration of drug action compared to a slow aqueous humor inlet. Case 2a and Case 2b at the middle vitreous provide the upper bounds for comparison for duration of action as the convection is 0 (slowest limiting cases for the corresponding boundary conditions). Case 1a with fast convection in rabbits keeps the concentration above the 2.6 µg/mL concentration threshold for 8.5 days compared to 18 days for slow aqueous humor inlet and 27.5 days for no aqueous humor inlet flow (no convection) and the same boundary conditions (Case 2a). In humans for Case 1a with fast convection, the duration of action is 11.5 days compared to 36 days with slow aqueous humor inlet and 67.5 days for no convection. For Case 1b in rabbits with fast convection, the concentration above the threshold lasts 2.5 days compared to 3 days when considering slow or no convection and in humans for 5 days compared to 8.5 days for slow convection and 9 days for no convection and the same boundary conditions (Case 2b).

## 4. Discussion

### 4.1. Effects of injection location on bevacizumab distribution after intravitreal injection

In rabbits, the concentration results for Case 1a and Case 2a are closer to the experimental data in all three injection locations (Figure 3a–c and Figure S7a–c). These results suggest that drug clearance in the rabbit eye is mainly driven by anterior elimination and that some limited convection might also play a role. Bevacizumab concentration in the rabbit vitreous drops rapidly when considering Case 1b and Case 2b, independent of the injection location, due to the posterior elimination. Even though experimental data is sparse, the data ^11–15^ still show a longer residence time in the vitreous than predicted by these two cases. This suggests posterior elimination is not the main route of drug clearance in rabbit vitreous, which is consistent with other published results ^41^.

In humans (Figure 3d–f and Figure S7d–f), Case 1b and Case 2b predict similar results, with these two cases being the closest to the experimental data at earlier time points, when the concentration is above the *in vivo* threshold concentration in the vitreous. This suggests that the posterior and the anterior elimination routes play significant roles in drug clearance from the human vitreous when concentrations at the fovea are high (Figure 4 and Figures S3–S6). Bevacizumab concentration in Case 1a is slightly closer to the experimental data at earlier time points than in Case 2a, but Case 1a is better at predicting later time points than the other cases, indicating that convection also likely plays a role, compared to Case 2a without convection. While none of the cases completely fits the limited data ^16,17^ exactly, the simulated cases do bound the actual human data. The later timepoint agreement with Case 1a suggests that the anterior elimination route is dominant at later times, likely due to the concentration at the fovea and near the posterior elimination boundary being depleted in the large human eye. Regardless of the injection location, Case 2a significantly overestimates drug concentration in the human vitreous. Together, these results indicate that the human boundary conditions are likely a combination of anterior elimination and posterior elimination, and the drug distribution includes some effects of convection.

The calculated half-lives (Table 3) agree with the PK profiles obtained in the rabbit vitreous and are closer to the lower end of the reported range of 3.51–7.06 days ^64^ when considering Case 2a in rabbits (Case 1a is also reasonable). The half-lives for the human vitreous are near the expected range of 5.8 ± 1.27 days ^65^, which matches Case 1a most closely. Another source that calculated vitreal elimination half-lives based on a retrospective literature survey has the bevacizumab half-lives as 5.5 days for rabbit and 10.1 days for human ^41^, which is most consistent with Case 2a calculated for rabbit and for human here. If a slower convection were considered, then the resulting half-life would be between that of Case 1a and Case 2a. If the human boundary conditions were to be adjusted to be a combination of anterior elimination and somewhat limited posterior elimination (the boundary condition *C* = 0 might be too extreme), then the resulting half-life could also be reasonable.

Bevacizumab half-life increases when injections are nearer to the posterior segment of the eye without posterior elimination. This is expected as the farther the injection site from the anterior segment, the slower the clearance of bevacizumab. In the larger human eye for cases where both anterior and posterior elimination are considered (Case 1b and Case 2b), injections at the anterior of the vitreous may provide marginally longer half-lives due to the relatively smaller surface area for anterior elimination proximal to the dose (Table 3).

The rapid initial increase in the concentration at the fovea for all cases (Figure 4) is due to the initial condition being zero and the reasonable rate of distribution throughout the entire vitreous within a few days (Figures S8–S11), even for the diffusion only cases. The sharp decrease characteristic of Case 1b and Case 2b is due to the way the concentration is calculated when posterior elimination is considered (see Section 2.6 for the procedure); the concentration is heavily influenced by the flux value. For all cases, the drug diffuses towards areas with lower concentration, and this effect is most pronounced at early times while the concentration gradient is large between the dose injection site and the rest of the domain. For a *C* = 0 boundary condition at the posterior segment, a concentration gradient exists at this boundary as long as some drug remains in the vitreous. This boundary condition maximizes the flux through the posterior elimination route. Injecting the drug closer to the posterior segment reduces the distance to the boundary and increases the local concentration and the difference between that concentration and the *C* = 0 boundary conditions, driving a larger flux (amount per area per time) to go through the posterior vitreous boundary. Faster flux results in a steeper concentration drop in the fovea.

As expected, calculated half-lives in the human vitreous were longer than the ones in the rabbit vitreous due to the size differences of the eyes (Table 3). The drug’s duration of action in the human vitreous is longer than in the rabbit vitreous (Table 4) since bevacizumab clearance in the smaller rabbit eye is faster than in the larger human eye. Note that only Case 1a for the human vitreous is within the expected duration of action interval of 28–42 days, which is usually the dosing interval for bevacizumab in humans. This is further consistent with the idea that the human conditions are likely a combination of anterior elimination and somewhat limited posterior elimination boundary conditions and convection, which would also generate an intermediate value of the duration of action between the values of the simulated cases.

### 4.2. Effects of convection on bevacizumab distribution after intravitreal injection in the middle vitreous

Khoobyar et al. ^38^ calculated a pressure distribution in the human eye considering the vitreous divided into a liquefied and a gel-type region. They calculated the pressure distribution throughout the vitreous and obtained a pressure drop below 50 Pa ^38^. However, our calculated pressure drop for low convection (Figure S13a) is consistent with other publications that report pressure drops below 2 Pa in the vitreous ^23,26,44^.

As shown in Figure S14, for Case 1a increasing convection results in a substantially faster bevacizumab clearance than for slow convection. For Case 1b, the increase in convection produces a relatively small increase in bevacizumab clearance from the eye. By considering a fast inlet of aqueous humor through the hyaloid membrane, the predicted bevacizumab concentrations deviate more from the experimental data in rabbit vitreous. This supports the idea that, in rabbit vitreous, convection effects are essentially negligible, and the anterior route of elimination is the dominant route. For Case 1a in humans, considering a fast inlet of aqueous humor through the hyaloid membrane results in bevacizumab concentrations closer to the experimental data in human vitreous for earlier timepoints. This supports the idea that vitreous convection can be present in humans, and both the anterior and posterior routes of elimination play significant roles in drug clearance.

In humans, the duration of action (Table 4) for Case 1a is within the expected range for frequency of administration (28–42 days) for slow convection. This suggests that convection in the human eye has velocities low compared to the overall aqueous humor production rate. Furthermore, considering adjusted boundary conditions as mentioned above may further improve the prediction of bevacizumab’s duration of action.

### 4.3. Current state of the art of existing related computational efforts

A 3D PK model for the rabbit eye was developed for bevacizumab distribution after both intravitreal injection and bevacizumab release from PLGA microspheres ^49,50^. Heat, momentum, and mass transport were considered in another 3D PK model after intravitreal injection of bevacizumab ^66^. The researchers considered diffusion and convection in the aqueous humor but only diffusion in the vitreous. Intravitreal injection was modeled as an instantaneous point source. The model’s boundary conditions included zero concentration at the iris and choroid. A limitation of that work is the model’s low accuracy in predicting bevacizumab concentration at early time points. Multiple computational investigations have simulated transport through the vitreous. The most common assumption is to consider the vitreous as a porous media and use Darcy’s law to solve the fluid flow problem ^21,23–25,38,52^. However, if a liquefied region is also considered within the vitreous, it is common to solve the fluid flow in porous media using Brinkman’s equation ^52,67^ to allow for convection due to shear stresses.

Our model results when considering anterior elimination only are either better or comparable at describing drug concentration in the vitreous after intravitreal injection in rabbits for the Bakri et al. ^11^ data when compared to previously published models of Sibak et al. ^66^ and Missel and Sarangapani ^68^ (Figure S16). One possible reason for the difference between the results of our rabbit model and published models (Figure S16) is that the published models ^66,68^ used bevacizumab diffusion coefficients of 4×10^*−*7^ and 5.21×10^*−*7^ cm^2^/s, respectively, which are smaller than our value of 1.2×10^*−*6^ cm^2^/s. However, an advantage of our model is the relative computational simplicity. Both these models involved an entire eye geometry, with a mesh of over 1.5 million elements in the model by Sibak et al. ^66^. In contrast, our rabbit model involved only the vitreous, with boundary conditions to account for neighboring eye regions, and a mesh of 162,750 elements.

Our model results when also considering posterior elimination better predict vitreous bevacizumab concentration after intravitreal injection in human eye for the Zhu et al. ^17^ data compared to the model by Missel and Sarangapani ^68^ (Figure S17), particularly at early time points. Later time points are better described when considering convection and anterior elimination only. The existing human model results ^68^ and our model results for the case of anterior elimination only are similar despite differences in the diffusion coefficients used (5.21×10^*−*7^ cm^2^/s in their model compared to 9.13×10^*−*7^ cm^2^/s in our model). Additionally, while their model involves a detailed human eye geometry, our model focuses specifically on the human vitreous, making the calculations less computationally expensive.

### 4.4. Limitations and model sensitivity

The finite element model used in the study is too computationally expensive for an exhaustive global sensitive analysis and formal uncertainty quantification. However, we systematically varied the important parameters of injection location, size of the vitreous (rabbit vs. human geometries), boundary conditions, and convective velocity in the results discussed above. The diffusion coefficient is an additional important parameter in the model. As we only considered one drug, bevacizumab, we did not show the impacts of varying the diffusion coefficient throughout this study. For a simple analysis of the impacts of changing the diffusion coefficient, we compare the half-life of bevacizumab in the human vitreous for Case 1a with slow convection and the dose injected in the middle vitreous position. With the value of *D* = 9.13 × 10^−11^ m^2^/s used throughout this study, the half-life is 5.36 days (Table 3). For a 50% increase in diffusion coefficient to 1.5*D*, the half-life reduces to 4.40 days (18% decrease). For increases to 2*D* and 3*D*, the half-lives reduce further to 3.69 days (31% decrease) and 2.73 days (50% decrease), respectively. The results are robust to small fluctuations in *D*, but large fold-changes expected for drugs of very different molecular weights do affect the results.

A limitation of the current work is that it focuses only on intravitreal injection. Others have considered modeling controlled-release formulations for intravitreal delivery (reviewed by Chacin Ruiz et al. ^9^). This is a future direction for extending the current work. Another limitation of our human model is the scarcity of available experimental data to better interpret and validate the results.

## 5. Conclusions

The drug transport mechanisms in the vitreous are not entirely understood and depend on several factors, including drug diffusivity, vitreous properties, boundary conditions, and individual ocular anatomy. These factors help explain the variations found in the experimental results where, under the same conditions, researchers obtained different PK profiles. Our 3D PK models successfully capture drug distribution in the vitreous of rabbits and humans and consider plausible elimination routes while comparing results to published data. Both our models’ results are close to the experimental data despite the high variability in the experimental data due to the difference in the vitreous’ physiological and material properties among individuals of the same species. Such simulation results can be used to gain insight into different dosing regimes.

Model results suggest that in rabbit vitreous, one of the most commonly used PK animal models, the effect of the anterior elimination route is dominant over the posterior elimination route. Differently, the results obtained for bevacizumab concentration in human vitreous suggest that the posterior elimination route plays a significant role alongside the anterior elimination route in drug clearance from the human eye. Furthermore, the modeling results show that drug injection location does not substantially influence drug half-life or duration of action in the vitreous, but the peak concentration experienced at the fovea is affected.

## Acknowledgments

This work was supported by National Institutes of Health grant R35GM133763 to ANFV, R01EB032870 to KESR, MPO, and ANFV, Owen Locke Foundation to KESR, and the University at Buffalo. We thank lab members and collaborators for their thorough feedback on this manuscript and helpful discussions.

