## Supplementary File for "Computer Modeling of Bevacizumab Drug Distribution After Intravitreal Injection in Rabbit and Human Eyes"

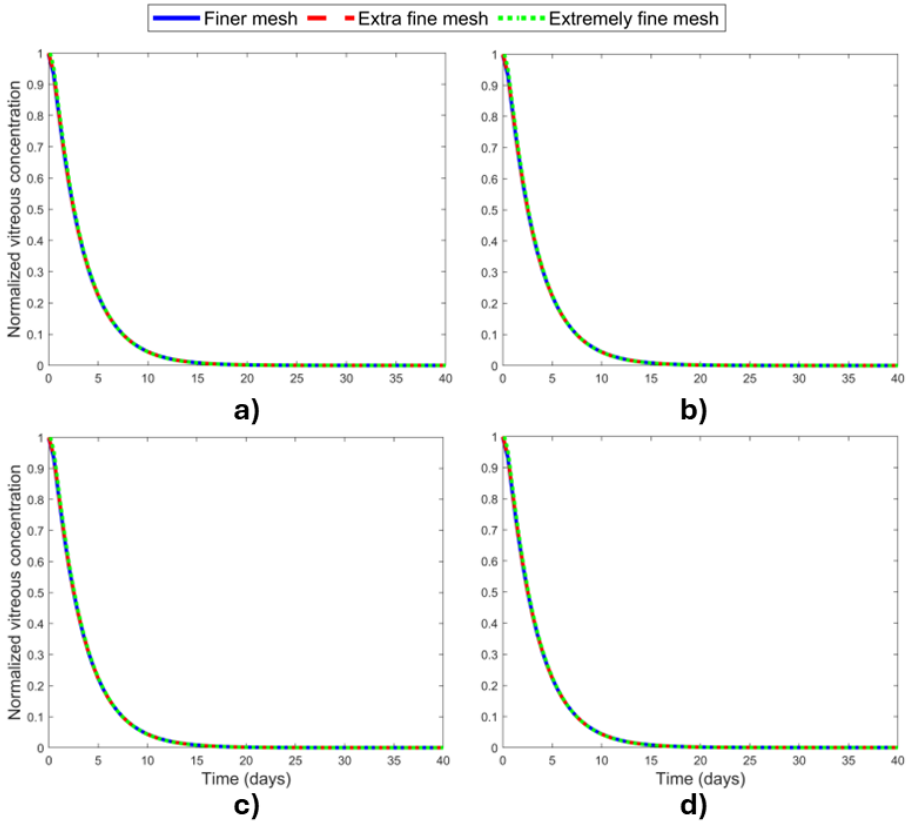


Figure S1: Mesh independence test for rabbit vitreous model comparing normalized vitreous concentration in the rabbit eye. Physics-defined sizes of “finer”, “extra fine”, and “extremely fine” meshes on COMSOL were compared for the dose injected in the middle vitreous area while considering slow convection. a) Case 1a: slow convective flow with anterior elimination only, b) Case 1b: slow convective flow with both anterior and posterior elimination, c) Case 2a: no convective flow with anterior elimination, and d) Case 2b: no convective flow with both anterior and posterior elimination.

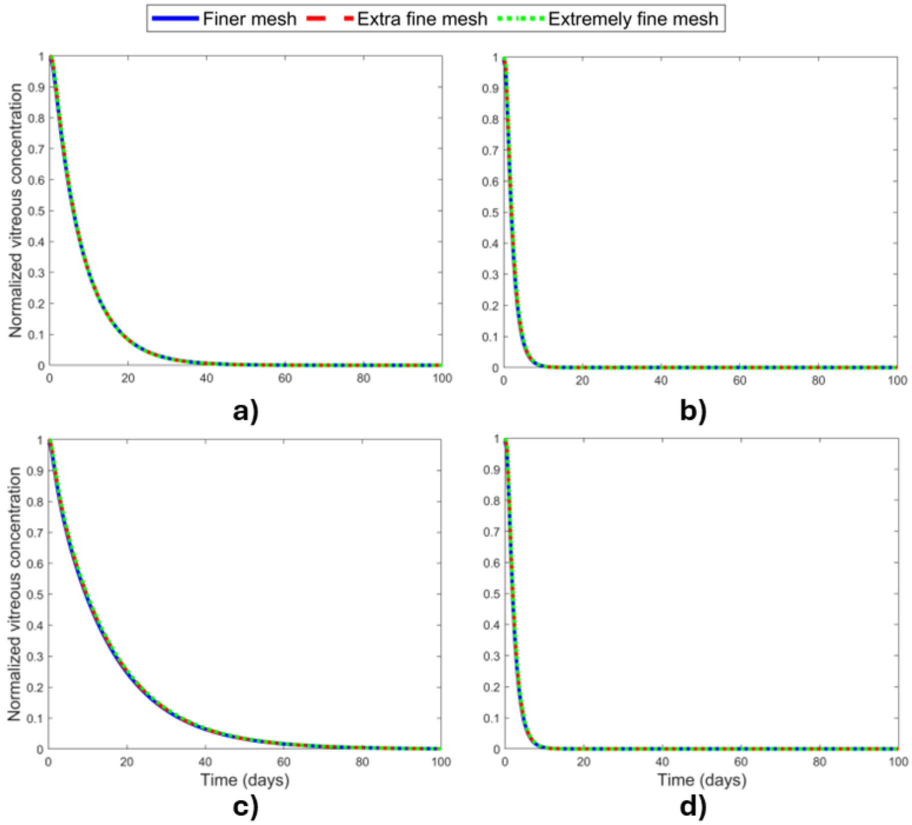


Figure S2: Mesh independence test for human vitreous model comparing normalized vitreous concentration in the human eye. Physics-defined sizes of “finer”, “extra fine”, and “extremely fine” meshes on COMSOL were compared for the dose injected in the middle vitreous area while considering slow convection. a) Case 1a: slow convective flow with anterior elimination only, b) Case 1b: slow convective flow with both anterior and posterior elimination, c) Case 2a: no convective flow with anterior elimination, and d) Case 2b: no convective flow with both anterior and posterior elimination.

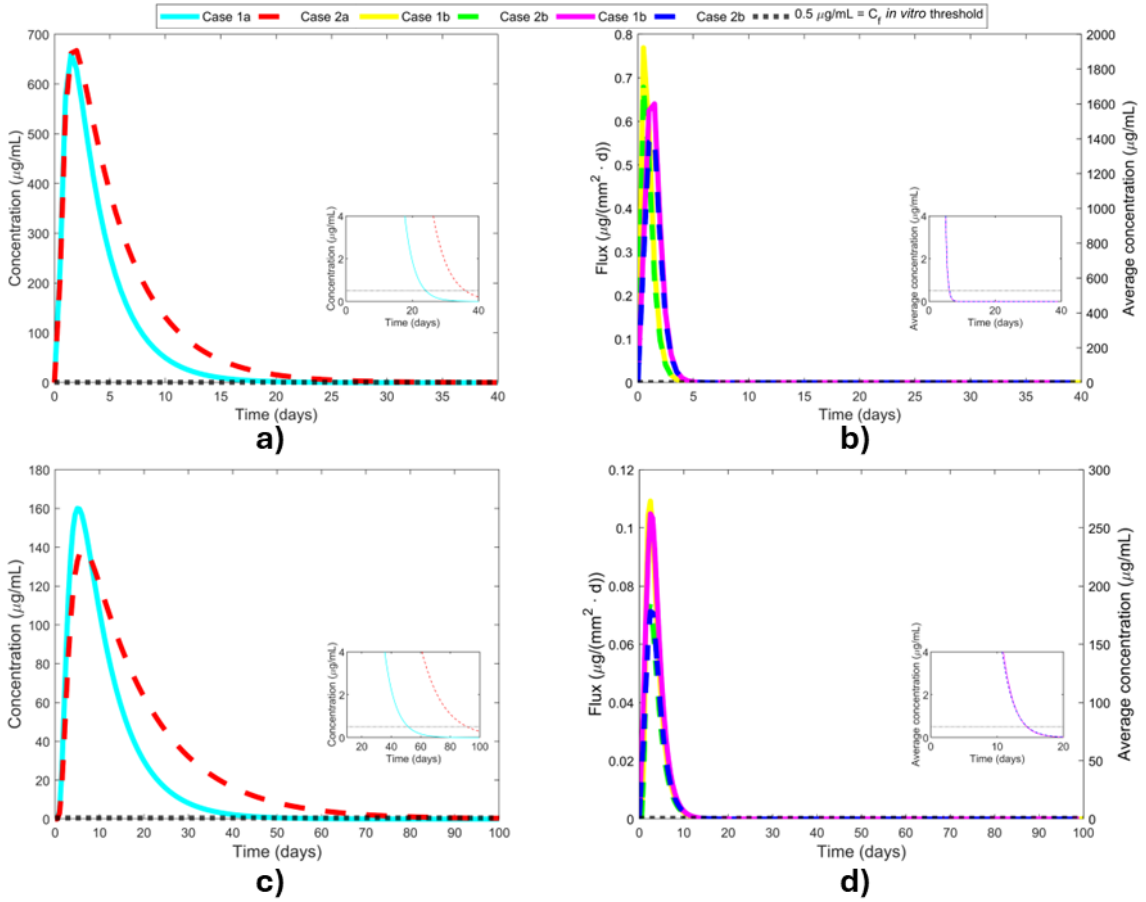


Figure S3: Drug concentration at the rabbit "fovea" and human fovea for the dose injected in the anterior vitreous considering slow convection. Concentrations a) directly measured from a point probe at the rabbit "fovea," b) calculated as the daily average based on flux measurements from a point probe and the spatial dimensions of the rabbit "fovea," c) directly measured from a point probe at the human fovea, and d) calculated as the daily average based on flux measurements from a point probe and the spatial dimensions of the human fovea. Case 1a: slow convective flow with anterior elimination only, Case 1b: slow convective flow with both anterior and posterior elimination, Case 2a: no convective flow with anterior elimination, and Case 2b: no convective flow with both anterior and posterior elimination. The yellow and green Case 1b and 2b curves are the flux values (left-side $y$-axis), and the pink and blue Case 1b and 2b curves are the average concentrations (right-side $y$-axis). $C_f$: *in vitro* threshold concentration at the fovea. Note that the $y$-axis scales are different for each subfigure.

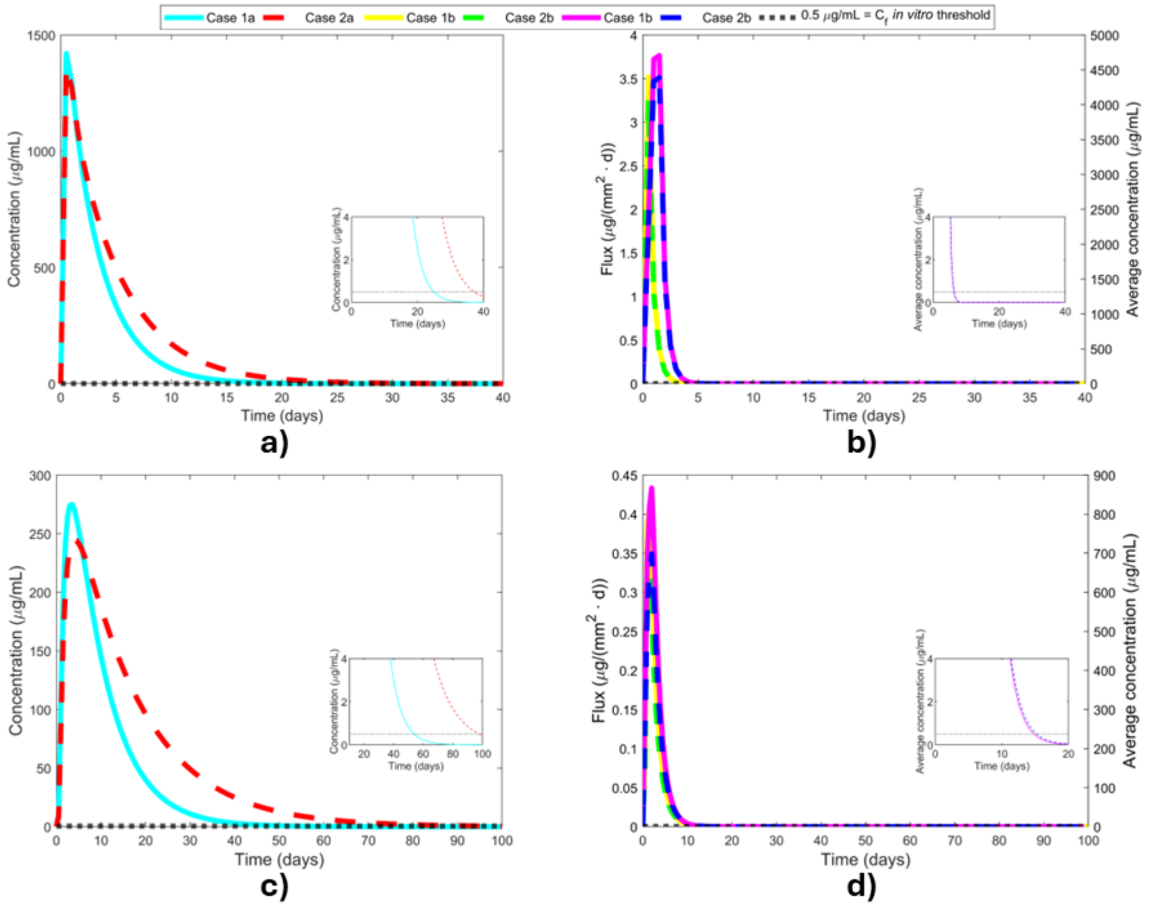


Figure S4: Drug concentration at the rabbit "fovea" and human fovea for the dose injected in the middle vitreous considering slow convection. Concentrations a) directly measured from a point probe at the rabbit "fovea," b) calculated as the daily average based on flux measurements from a point probe and the spatial dimensions of the rabbit "fovea," c) directly measured from a point probe at the human fovea, and d) calculated as the daily average based on flux measurements from a point probe and the spatial dimensions of the human fovea. Case 1a: slow convective flow with anterior elimination only, Case 1b: slow convective flow with both anterior and posterior elimination, Case 2a: no convective flow with anterior elimination, and Case 2b: no convective flow with both anterior and posterior elimination. The yellow and green Case 1b and 2b curves are the flux values (left-side $y$-axis), and the pink and blue Case 1b and 2b curves are the average concentrations (right-side $y$-axis). $C_f$: *in vitro* threshold concentration at the fovea. Note that the $y$-axis scales are different for each subfigure.

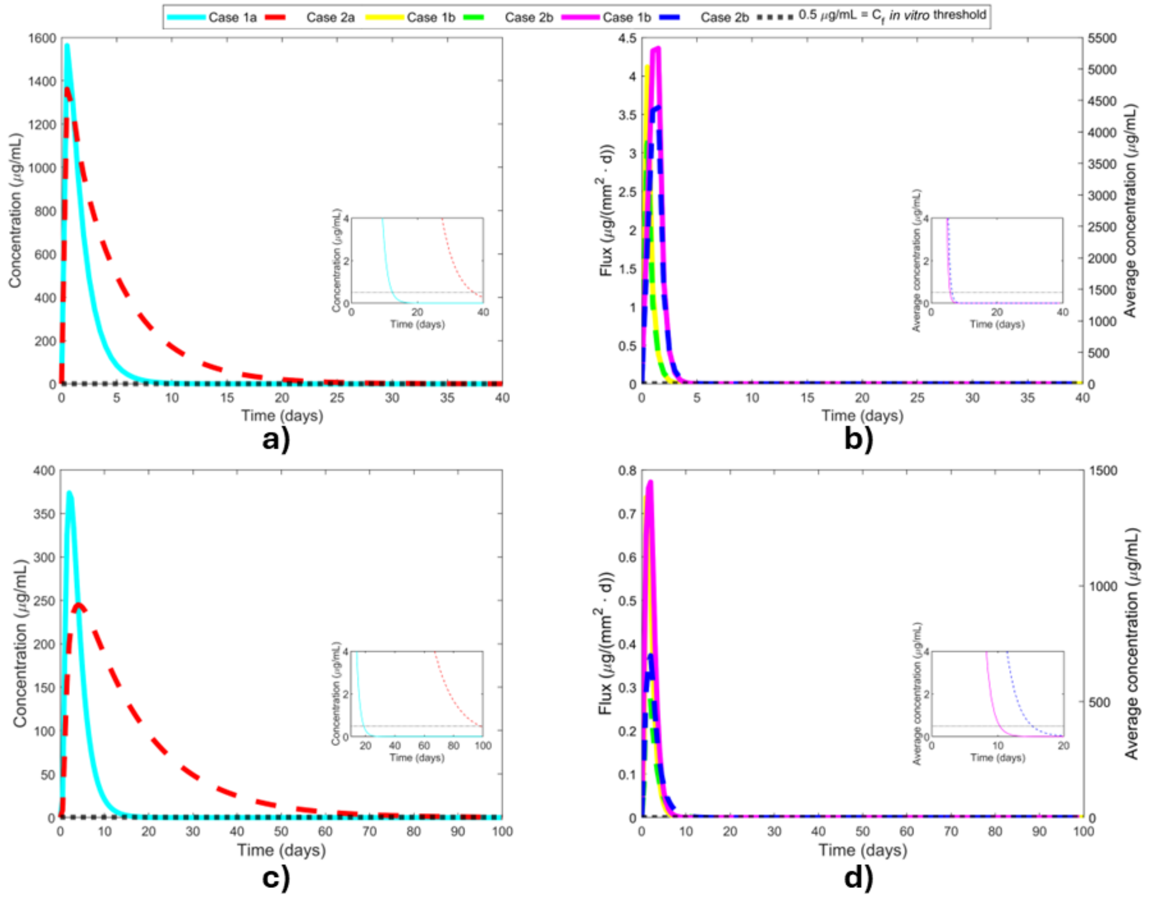


Figure S5: Drug concentration at the rabbit "fovea" and human fovea for the dose injected in the middle vitreous considering fast convection. Concentrations a) directly measured from a point probe at the rabbit "fovea," b) calculated as the daily average based on flux measurements from a point probe and the spatial dimensions of the rabbit "fovea," c) directly measured from a point probe at the human fovea, and d) calculated as the daily average based on flux measurements from a point probe and the spatial dimensions of the human fovea. Case 1a: fast convective flow with anterior elimination only, Case 1b: fast convective flow with both anterior and posterior elimination, Case 2a: no convective flow with anterior elimination, and Case 2b: no convective flow with both anterior and posterior elimination. The yellow and green Case 1b and 2b curves are the flux values (left-side $y$-axis), and the pink and blue Case 1b and 2b curves are the average concentrations (right-side $y$-axis). $C_f$: *in vitro* threshold concentration at the fovea. Note that the $y$-axis scales are different for each subfigure.

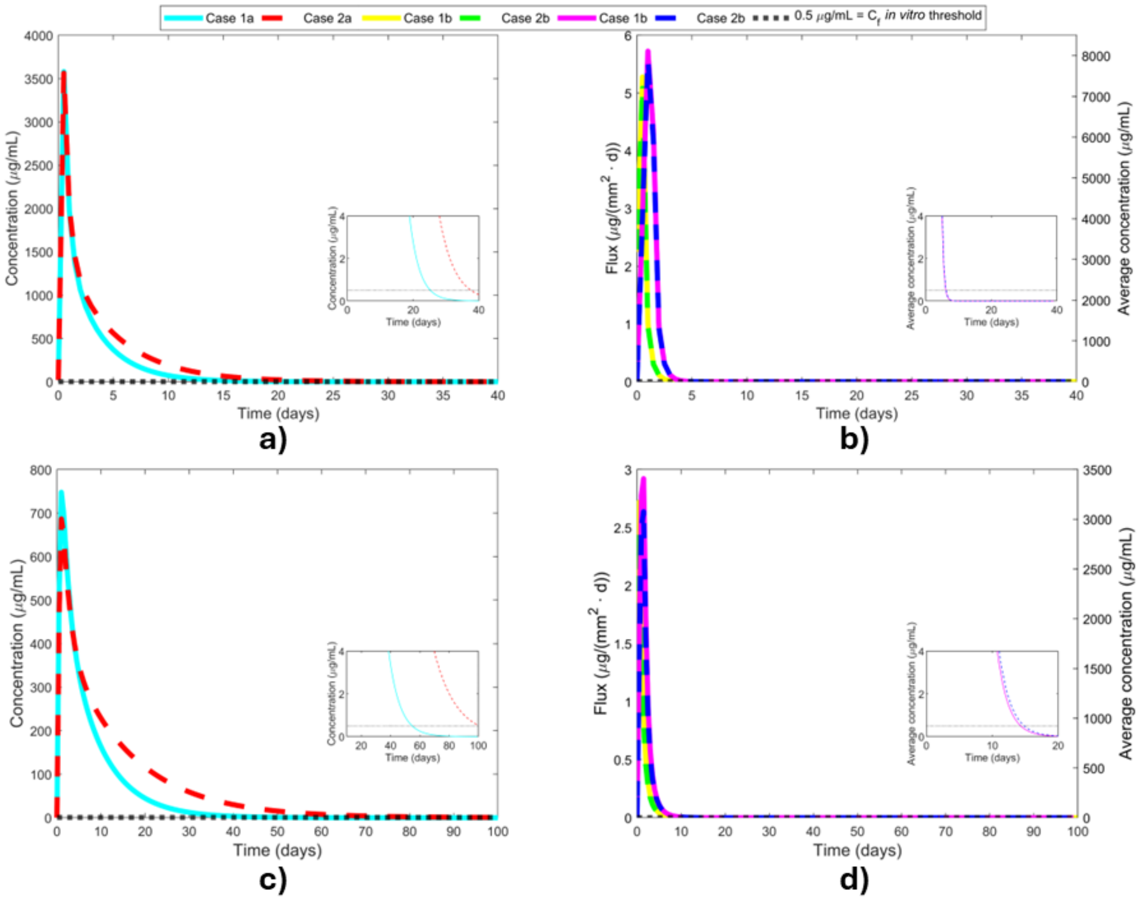


Figure S6: Drug concentration at the rabbit "fovea" and human fovea for the dose injected in the posterior vitreous considering slow convection. Concentrations a) directly measured from a point probe at the rabbit "fovea," b) calculated as the daily average based on flux measurements from a point probe and the spatial dimensions of the rabbit "fovea," c) directly measured from a point probe at the human fovea, and d) calculated as the daily average based on flux measurements from a point probe and the spatial dimensions of the human fovea. Case 1a: slow convective flow with anterior elimination only, Case 1b: slow convective flow with both anterior and posterior elimination, Case 2a: no convective flow with anterior elimination, and Case 2b: no convective flow with both anterior and posterior elimination. The yellow and green Case 1b and 2b curves are the flux values (left-side $y$-axis), and the pink and blue Case 1b and 2b curves are the average concentrations (right-side $y$-axis). $C_f$: *in vitro* threshold concentration at the fovea. Note that the $y$-axis scales are different for each subfigure.

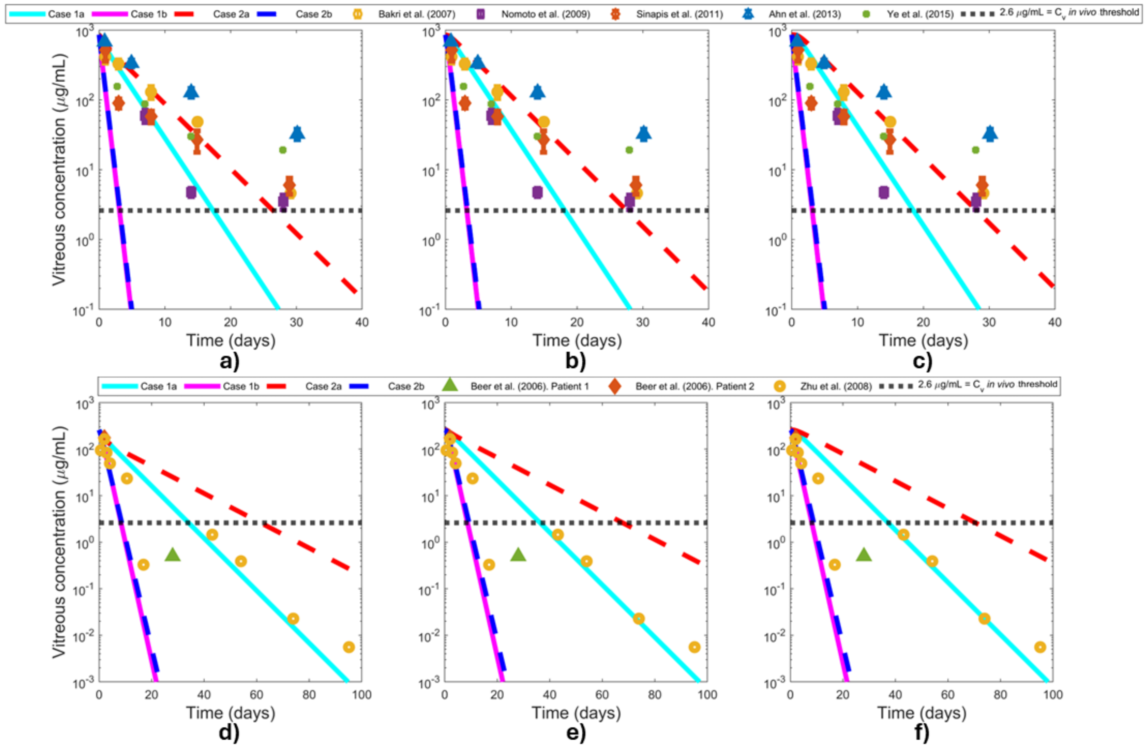


Figure S7: Logarithmic vitreous concentration in the rabbit and human eye. Injection locations: a) rabbit anterior vitreous, b) rabbit middle vitreous, c) rabbit posterior vitreous, d) human anterior vitreous, e) human middle vitreous, and f) human posterior vitreous. Case 1a: slow convective flow with anterior elimination only, Case 1b: slow convective flow with both anterior and posterior elimination, Case 2a: no convective flow with anterior elimination, and Case 2b: no convective flow with both anterior and posterior elimination. Model results for each case are shown as curves. Experimental data (markers) from Bakri et al.[11], Nomoto et al.[12], Sinapis et al.[13], Ahn et al.[14], Ye et al.[15], Beer et al.[16], and Zhu et al.[17]. $C_V$: *in vivo* threshold concentration in the vitreous.

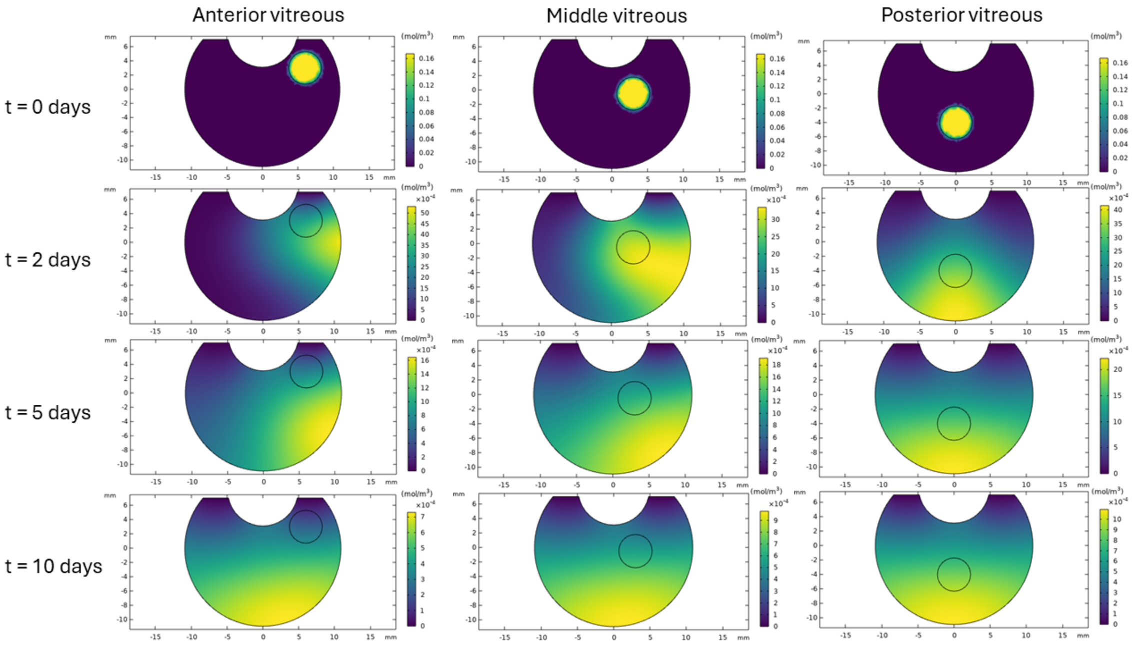


Figure S8: Bevacizumab concentration (mol/m$^3$) at the human vitreous. Slow convective flow with anterior elimination only (Case 1a). Note that the scale bars are different for each subfigure.

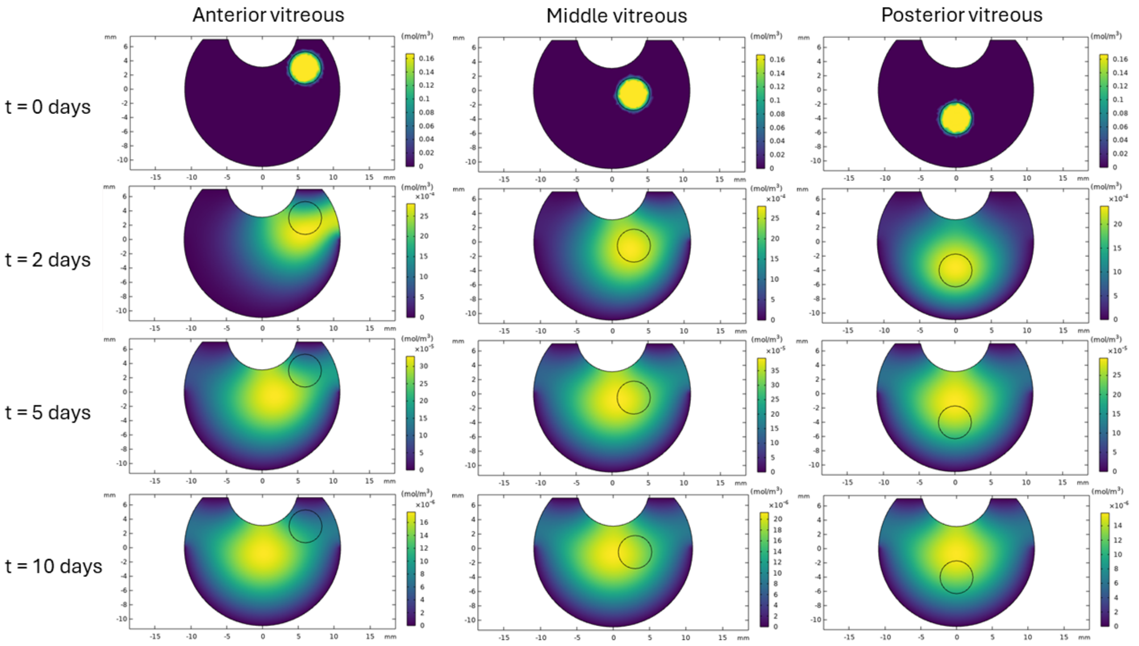


Figure S9: Bevacizumab concentration (mol/m$^3$) at the human vitreous. Slow convective flow with both anterior and posterior elimination (Case 1b). Note that the scale bars are different for each subfigure.

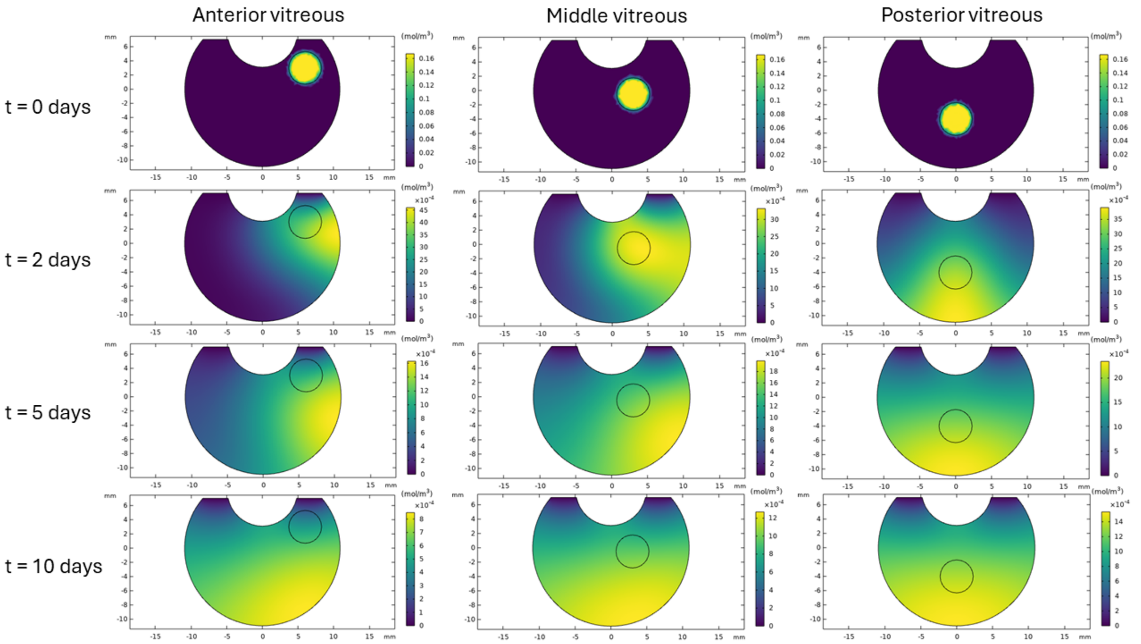


Figure S10: Bevacizumab concentration (mol/m$^3$) at the human vitreous. No convective flow with anterior elimination (Case 2a). Note that the scale bars are different for each subfigure.

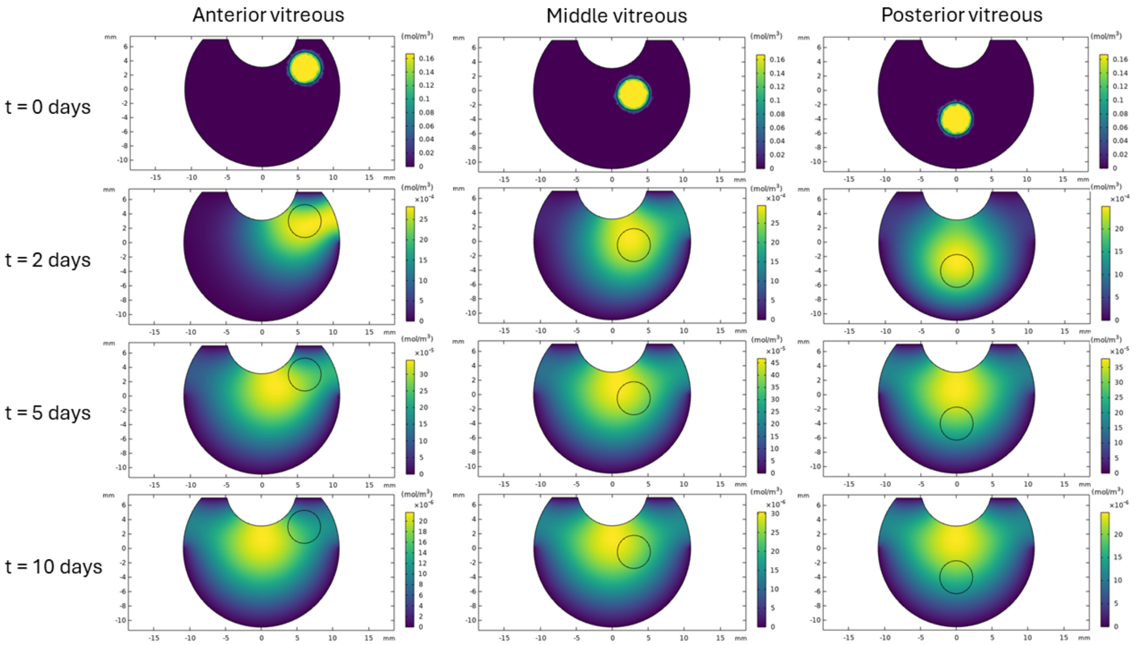


Figure S11: Bevacizumab concentration (mol/m$^3$) at the human vitreous. No convective flow with both anterior and posterior elimination (Case 2b). Note that the scale bars are different for each subfigure.

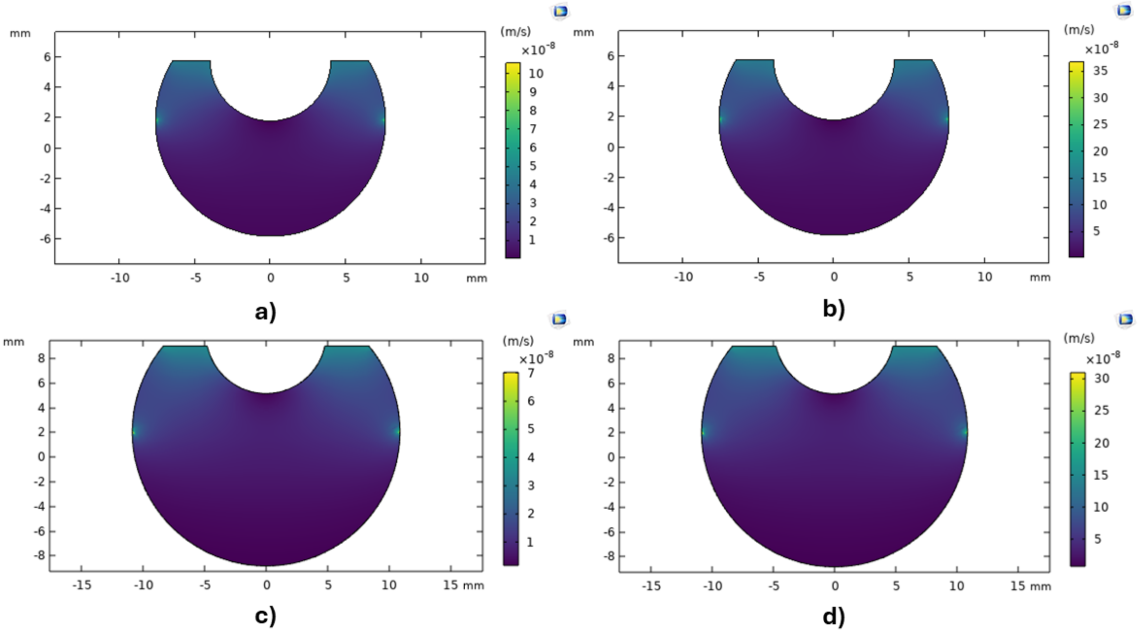

Figure S12: Velocity (m/s) profiles at the center of the rabbit and human vitreous at steady state for a) slow inlet at rabbit hyaloid membrane ($4.3 \times 10^{-8}$ m/s), b) fast inlet at rabbit hyaloid membrane ($1.5 \times 10^{-7}$ m/s), c) slow inlet at human hyaloid membrane ($3.4 \times 10^{-8}$ m/s), and d) fast inlet at human hyaloid membrane ($1.5 \times 10^{-7}$ m/s). Note that the scale bars are different for each subfigure. The phenomena are the same, but the magnitude of the effect is scaled relative to the velocity and geometry.

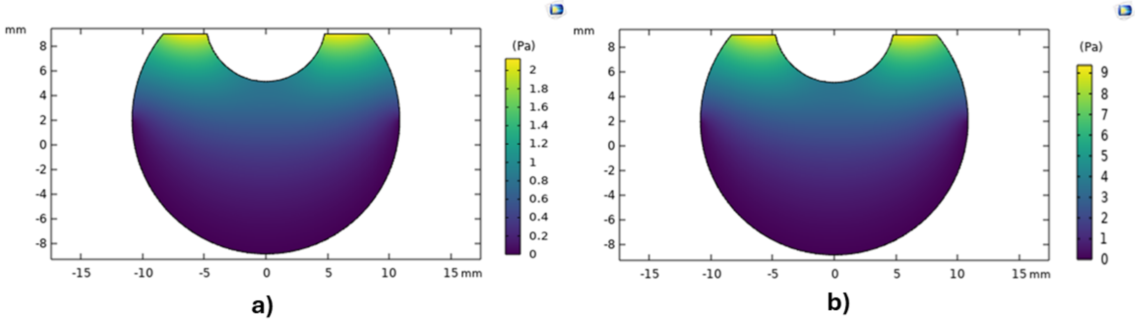

Figure S13: Pressure (Pa) profile in human vitreous at steady state for a) slow convection and b) fast convection. Note that the scale bars are different for each subfigure. The phenomena are the same, but the magnitude of the effect is scaled relative to the velocity.

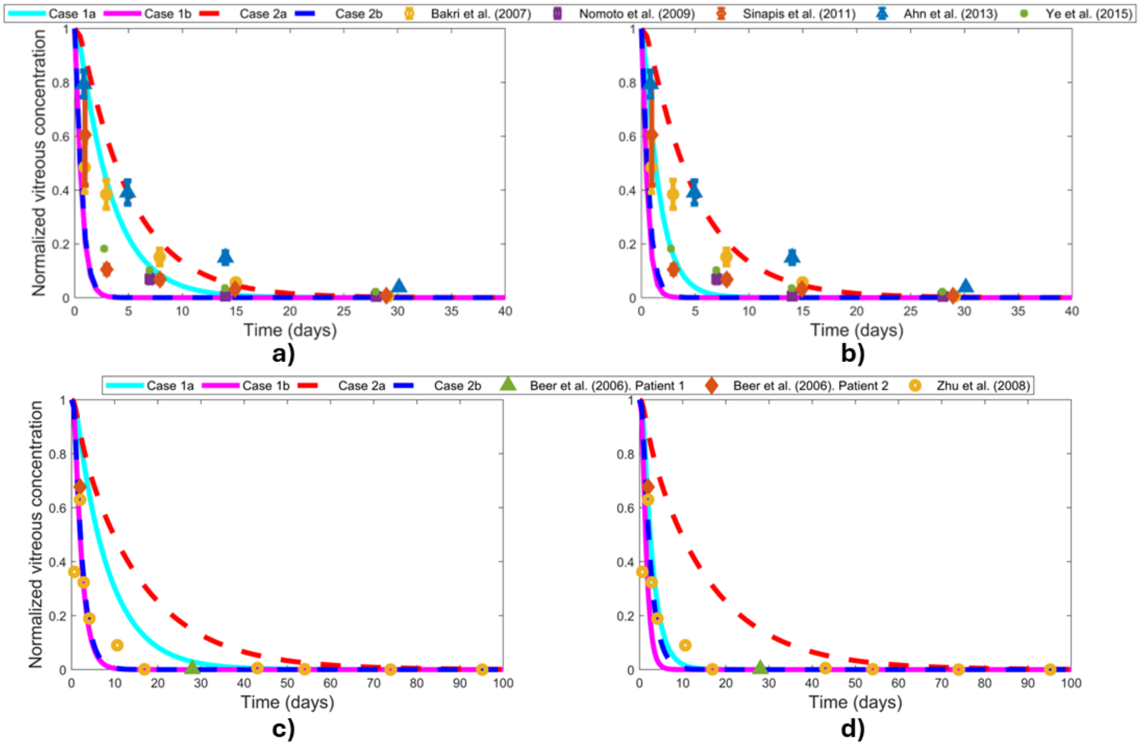


Figure S14: Normalized vitreous concentration in the rabbit and human eye for the dose injected in the middle vitreous. Scenarios: a) slow convection in rabbit, b) fast convection in rabbit, c) slow convection in human, and d) fast convection in human. Case 1a: convective flow with anterior elimination only, Case 1b: convective flow with both anterior and posterior elimination, Case 2a: no convective flow with anterior elimination, and Case 2b: no convective flow with both anterior and posterior elimination. Model results for each case are shown as curves. Experimental data (markers) are from Bakri et al.[11], Nomoto et al.[12], Sinapis et al.[13], Ahn et al.[14], Ye et al.[15], Beer et al.[16], and Zhu et al.[17].

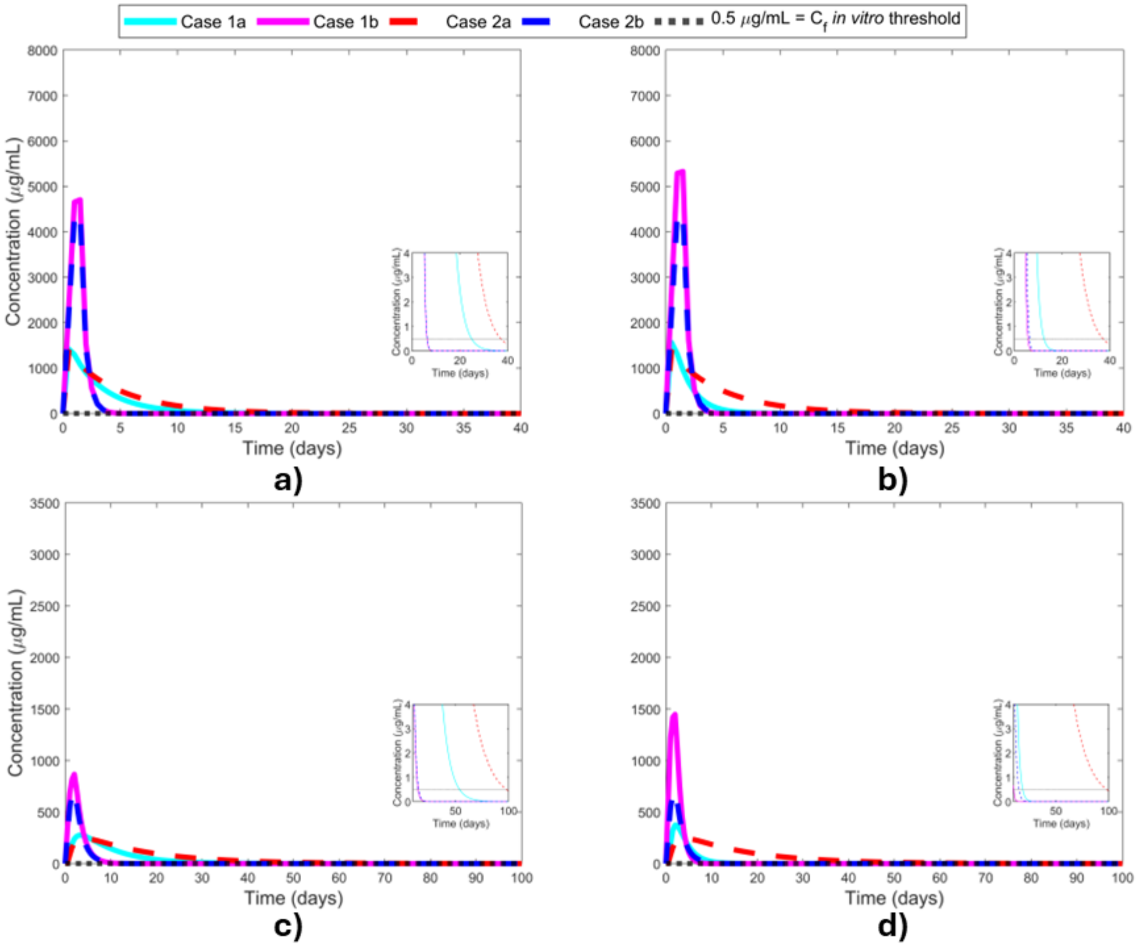


Figure S15: Drug concentration at the rabbit "fovea" and human fovea for the dose injected in the middle vitreous. Scenarios: a) slow convection in rabbit, b) fast convection in rabbit, c) slow convection in human, and d) fast convection in human. Case 1a: convective flow with anterior elimination only, Case 1b: convective flow with both anterior and posterior elimination, Case 2a: no convective flow with anterior elimination, and Case 2b: no convective flow with both anterior and posterior elimination. $C_f$: *in vitro* threshold concentration at the fovea.

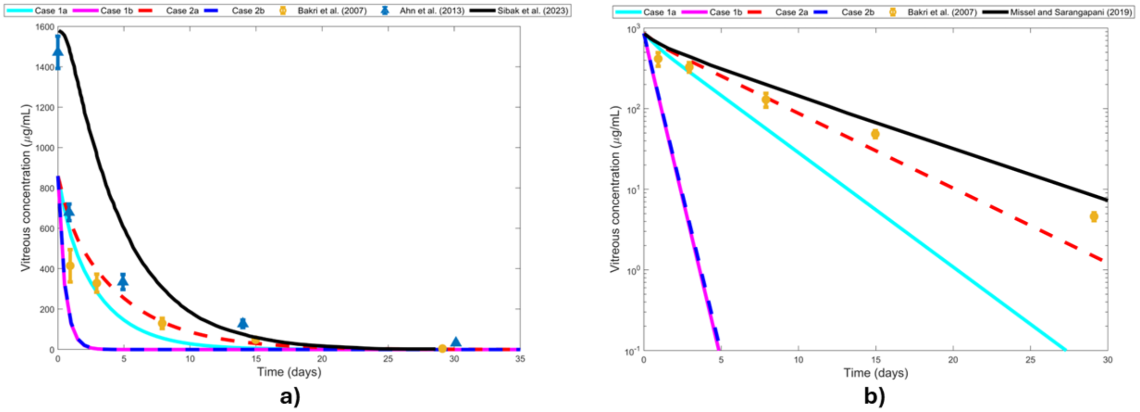


Figure S16: Comparison of the rabbit model against existing rabbit ocular models: a) model by Sibak et al.[66] and b) model by Missel and Sarangapani[68]. We consider the anterior vitreous as the injection location since it is the location used in both published models. Case 1a: slow convective flow with anterior elimination only, Case 1b: slow convective flow with both anterior and posterior elimination, Case 2a: no convective flow with anterior elimination, and Case 2b: no convective flow with both anterior and posterior elimination. Our model results for each case are shown as colored curves. Existing model results are shown as black curves. WebPlotDigitizer[62] was used to obtain the values for these curves. Experimental data (markers) from Bakri et al.[11] and Ahn et al.[14].

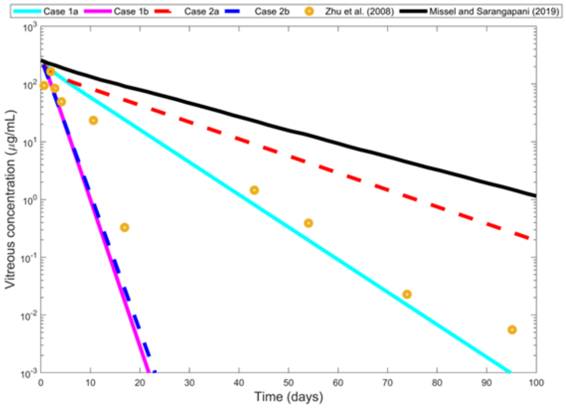


Figure S17: Comparison of our human model against the existing human ocular models by Missel and Sarangapani[68]. We consider the anterior vitreous as the injection location since it is the location used in this published model. Case 1a: slow convective flow with anterior elimination only, Case 1b: slow convective flow with both anterior and posterior elimination, Case 2a: no convective flow with anterior elimination, and Case 2b: no convective flow with both anterior and posterior elimination. Our model results for each case are shown as colored curves. Existing model results are shown as black curves. WebPlotDigitizer[62] was used to obtain the values for these curves. Experimental data (markers) from Zhu et al.[17].
